# Single-cell spatiotemporal transcriptomics reveals the origin, cellular heterogeneity and regulatory network of seed fiber cells in poplar

**DOI:** 10.64898/2026.03.28.714976

**Authors:** Kaiyuan Han, Hanxin Wang, Jingyun Tan, Xiong Yang, Tianyun Zhao, Liming Jia, Xinmin An, Zhong Chen

## Abstract

Poplar (*Populus* spp.) is globally important for timber and afforestation, and is also a model system for woody perennials. However, the molecular control of reproductive organ development, particularly that of the epidermal trichomes on seeds known as seed fibers (fluff), which cause environmental and health concerns, remains poorly understood. Here, integrating single-nucleus RNA sequencing (snRNA-seq) with spatial transcriptomics, we constructed a spatiotemporal atlas of female poplar capsules that resolves the cellular composition of the organ and positions each cell type within its native tissue context. We traced the developmental trajectory of seed fibers, showing that they originate from placenta epidermal cells, and identified three functionally distinct fiber cell subtypes involved in initiation, photosynthesis and elongation. Network analysis identified candidate regulators including *PtoPDF2*, *PtoMYB38* and *PtoHDG11*. PtoPDF2 activates *PtoMYB38* and *PtoSTK2* by binding L1-box motifs in their promoters, revealing a module that governs seed fiber initiation. Together, our study provides a framework for seed fiber development in poplar and candidate targets for breeding low-fluff cultivars.

**Highlights:**

1. Single-cell spatiotemporal atlas of poplar capsule development
2. Trajectory analysis traces fiber cells to the placenta epidermis and identifies three distinct subtypes
3. PtoPDF2–*PtoMYB38*/*PtoSTK2* defines a new regulatory module driving seed fiber formation in poplar

## Introduction

Poplar (*Populus* spp.) trees are widely distributed across the Northern Hemisphere and serve as one of the most important afforestation species, playing critical roles in timber production, ecological restoration, and carbon sequestration^1–3^. However, the annual springtime release of massive seed fibers (fluff, seed trichomes, or catkins) from female trees has become a persistent environmental and public health concern in urban areas, causing air pollution, respiratory diseases, and fire hazards. These fibers develop inside the capsules of female trees and are released upon capsule dehiscence. Current management strategies relying on physical interventions and chemical treatments are costly, labor-intensive, and pose ecological risks^4^. One strategy to mitigate this problem has been to avoid planting female trees through early sex identification. Recent studies have identified *ARABIDOPSIS RESPONSE REGULATOR 17* (*ARR17*^5^) in *Populus tremula* and *Female-specifically expressed RESPONSE REGULATOR* (*FERR*^6^) in *Populus deltoides* as key genes that promote female development, and in males of *P. deltoides* a Y-chromosome-specific copy of FERR produces small interfering RNAs that suppress the functional *FERR* gene, thereby directing male development^6^. These findings enable sex identification at the seedling stage but address sex differentiation rather than the morphogenesis of seed fibers themselves. As the direct structural source of fluff, the cellular differentiation and molecular mechanisms governing seed fiber development remain poorly understood, leaving genetic improvement without precise targets.

Beyond its applied significance, poplar serves as a premier model system for woody perennial biology, with a well-annotated genome and extensive genomic resources^7,8^. However, compared to the substantial progress in understanding vegetative development, the molecular control of reproductive organ development, particularly the cellular differentiation of seed fibers, remains poorly characterized. Seed fibers are epidermal trichomes that arise from the placenta^4^, representing a unique system for studying reproductive organ trichome development in woody perennials. Elucidating the developmental logic of seed fiber formation therefore not only addresses a practical need for fluff mitigation but also provides fundamental insights into the evolution and regulation of trichome development in woody perennials, a largely unexplored frontier in plant developmental biology.

Plant trichomes exhibit remarkable diversity in morphology, cellular composition, and developmental origin across species. In *Arabidopsis thaliana*, unicellular trichome development depends on a conserved R2R3MYB-bHLH-WD40 (MBW) transcriptional activator complex comprising GLABRA1 (GL1/MYB), GLABRA3 (GL3/bHLH), and TRANSPARENT TESTA GLABRA1 (TTG1/WD40)^9^. This complex activates downstream genes including *GLABRA2* (*GL2*) to promote epidermal cell differentiation^10^, and its activity is modulated by plant hormones such as gibberellins and jasmonates^11,12^, as well as epigenetic regulators including chromatin assembly factors and histone acetyltransferases^13,14^. Cotton (*Gossypium hirsutum*) fibers, also unicellular trichomes, originate from ovule epidermal cells and rely on distinct regulators including *GhMYB2*^15^, *GhMYB25-like*, and *Homeobox-leucine zipper protein HOX3* (*GhHOX3*^16^), with notable differences from the *Arabidopsis* MBW pathway (Tian et al., 2020). Multicellular trichomes in tomato (*Solanum lycopersicum*) and cucumber (*Cucumis sativus*) exhibit even greater complexity, involving cell cycle regulators such as *Cyclin B2* (*SlCycB2*^17^) and intricate hormone signaling networks integrating jasmonate and auxin pathways^18,19^.

In poplar, studies on leaf trichomes have begun to reveal key regulatory genes and mechanisms. Activation tagging of *PtaMYB186*, an ortholog of *Arabidopsis MYB106*, produced the fuzzy mutant with increased foliar trichome density, enhanced growth, photosynthesis, and insect resistance^20^. Additionally, the *miR319a/TCP* module was shown to act synergistically with gibberellin (GA) signaling to regulate leaf trichome initiation in *P. tomentosa*, with TEOSINTE BRANCHED/CYCLOIDEA/PCF 19 (TCP19) interacting with REPRESSOR OF ga1-3 (RGA) to repress GL1-induced *GL2* expression^21^. These findings suggest that a genetic framework for leaf trichome development in poplar is emerging, involving MYB transcription factors, *miR319/TCP* modules, and GA signaling.

In contrast, trichomes in reproductive organs, such as poplar seed fibers, have received far less attention. Poplar seed fibers superficially resemble cotton fibers but exhibit a unique developmental origin, arising from placenta epidermal cells rather than ovule integuments^4^. To date, only a limited number of genes have been implicated in poplar seed fiber formation. RNAi-mediated suppression of *AGAMOUS* and *SEEDSTICK* homologs, including *STK1* and *STK2*, reduced seed hair production but compromised seed viability^22^. CRISPR-based knockout of *CENTRORADIALIS* and *MYB* alleles, including *MYB38*, *MYB138*, and *MYB186*, generated fluffless capsules^23^. Ectopic expression of sucrose metabolism-related genes *sucrose synthase 2 (PtoSUS2*) and *vacuolar invertase 3* (*PtoVIN3*) increased trichome numbers in *Arabidopsis*^4^. In addition, the R2R3-MYB transcription factor PdeMIXTA04 was shown to promote trichome density upon overexpression in *Arabidopsis* and to interact with PdeMYC and PdeWD40^24^. However, these studies represent isolated gene discoveries rather than systematic dissection of the regulatory networks governing seed fiber development at cellular resolution.

Single-cell sequencing technologies offer unprecedented opportunities to dissect developmental processes at cellular resolution. Conventional bulk RNA-seq averages transcriptional signals across heterogeneous cell populations, masking critical cell type-specific dynamics during poplar capsule development^25^. Single-nucleus RNA sequencing (snRNA-seq) has emerged as a robust alternative for woody plants, circumventing cell wall removal challenges and minimizing stress-related artifacts^26,27^. This approach has proven powerful in resolving cell type-specific transcriptional programs in poplar vegetative organs, including stems^28^ and shoot apices^29^. In lignified stems, for instance, snRNA-seq captured previously underrepresented cell types such as vessel-associated cells and identified *MYB48* as a key regulator of their function^30^. However, snRNA-seq alone loses spatial context during tissue dissociation. Spatial transcriptomics addresses this limitation by preserving tissue architecture while capturing gene expression information, enabling precise mapping of cell types back to their original positions and revealing intercellular communication networks, as demonstrated in studies of *Arabidopsis* leaf development^31^, rice (*Oryza sativa*) root zonation^32^, cotton fiber initiation^33^, and Norway spruce reproductive development^34^. However, the application of this integrated approach at single-cell resolution to seed fiber development in poplar has not yet been explored.

In this study, we integrated snRNA-seq and spatial transcriptomics to construct a high-resolution spatiotemporal atlas of female poplar capsule development. This integrative framework resolved the cellular composition of the developing capsule, mapped the spatial organization of distinct cell types within the tissue architecture, and traced the origin of seed fibers to placenta cells. Through single-cell pseudotemporal analysis, we reconstructed the developmental trajectory of fiber cells and identified three transcriptionally distinct subtypes associated with initiation, light responsiveness, and elongation. Weighted gene co-expression network analysis pinpointed multiple candidate regulators within the initiation-associated module, including *PROTODERMAL FACTOR 2* (*PtoPDF2*), *PtoMYB38*, and *HOMEODOMAIN GLABROUS 11* (*PtoHDG11*). Mechanistically, we demonstrated that PtoPDF2 activates the expression of *PtoMYB38* and *PtoSTK2* through direct binding to L1-box motifs in their promoters, unveiling a new transcriptional module that may govern seed fiber formation. Collectively, this integrative framework provides a systematic view of the cellular and molecular programs underlying seed fiber development in poplar and delivers a spatially resolved resource for dissecting cell fate decisions in woody perennial reproductive organs, offering candidate molecular targets for breeding low-fluff cultivars.

## Results

### Single-cell transcriptomic landscape of poplar capsule development

To elucidate the morphological dynamics of poplar seed fiber development, which initiates and matures rapidly within days post-anthesis (DPA)^4^, we conducted high-resolution cytological observations at 8-hour intervals starting from 3 DPA (**Supplementary Fig. 1**). During this period, the stigma at the capsule apex transitioned sequentially from pink to white and eventually to brown. Concurrently, the capsule wall shifted from yellow to dark green, accompanied by a gradual increase in capsule volume. At 4 DPA, pronounced nuclear staining was observed in the epidermal cells of the placenta, marking the initiation stage. By 5 DPA, these epidermal cells had differentiated into fiber cells, with visible protrusions marking the protrusion stage. At 6 DPA, these fiber cells underwent rapid elongation, progressing toward maturation, representing the elongation stage. Therefore, we collected samples at three key stages of seed fiber development: initiation (4 DPA 24:00), protrusion (5 DPA 24:00), and elongation (6 DPA 16:00), with three biological replicates per stage, and performed snRNA-seq using the 10x Genomics platform (**Fig. 1A**). High correlations were observed among the three biological replicates within each stage (**Supplementary Fig. 2**), demonstrating the reproducibility of our experimental design. After quality control and filtering at both the cellular and gene levels (**Supplementary Fig. 3**), we captured 7,741 to 12,611 cells from the nine samples across the three stages. The median genes per cell ranged from 1,180 to 1,761, and the median UMI counts per cell ranged from 1,773 to 2,785 (**Supplementary Table 1**). These metrics indicate the high quality of the dataset, making it suitable for subsequent analyses.

**Fig. 1.**
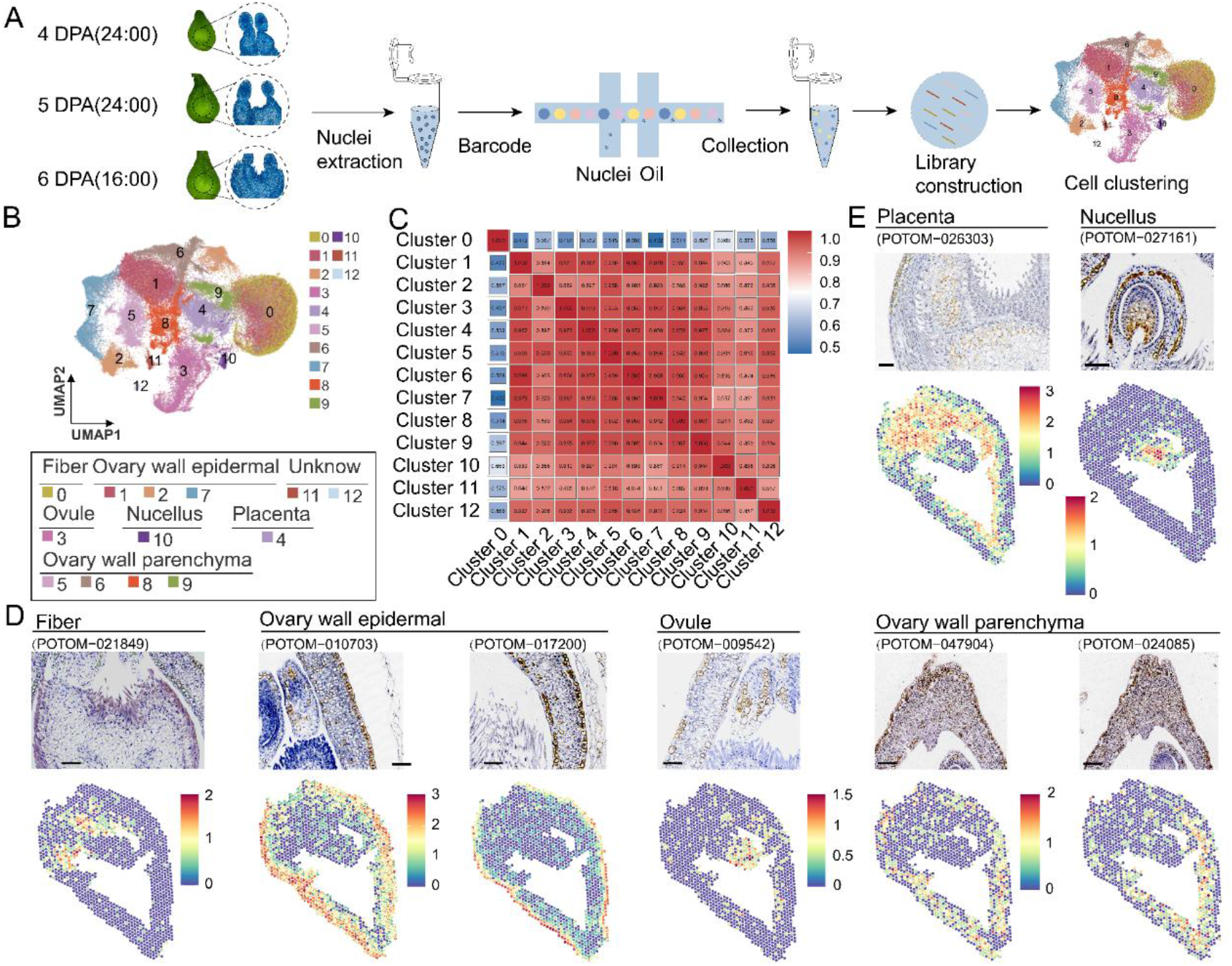
| Single-cell transcriptomic atlas of poplar female capsules. (**A**) Schematic overview of the experimental workflow for single-nucleus RNA sequencing of poplar capsules collected at three developmental stages. (**B**) Upper panel: UMAP visualization of unsupervised clustering results from all samples. Each dot represents a single cell. Lower panel: Identification of cell clusters, revealing six distinct cell types. (**C**) Heatmap showing the correlation coefficients among cell clusters. (**D**) Spatial expression patterns and RNA in situ hybridization validation of marker genes for fiber cells, ovary wall outer epidermal cells, ovary wall parenchyma cells, and ovule cells. (**E**) Spatial expression patterns and RNA in situ hybridization validation of marker genes for placenta cells and nucellus cells. Scale bars, 50 μm. Each gene was detected in at least three independent biological replicates from separate plants.

Dimensionality reduction and clustering analysis using the Uniform Manifold Approximation and Projection (UMAP) algorithm revealed a total of 13 distinct cell clusters (**Fig. 1B**). Correlation assessment of transcriptional profiles among these clusters showed that cluster 0 exhibited relatively low correlations with other clusters, suggesting that it may represent a functionally specialized cell population (**Fig. 1C**). Given the high tissue specificity of poplar capsules, we further performed spatial transcriptome sequencing on synchronous samples from the second stage (5 DPA 24:00) to obtain spatial expression information of genes. After rigorous initial quality assessment of the raw data and evaluation of nGene (number of genes), nUMI (number of UMIs), and percent.mt (mitochondrial gene percentage) per spot across samples, the data were confirmed to be of high quality (**Supplementary Table 2 and Fig. 4**). Subsequently, we compared the number of spots and median genes at different resolutions using BSTMatrix and found that the clustering results at level 6 (L6) provided a more accurate visualization that better matched the cellular characteristics of poplar capsules (**Supplementary Table 3 and Fig. 5**). To integrate the single-cell and spatial transcriptome datasets, we first performed Multimodal Interaction Analysis^35^ to evaluate the enrichment scores of marker gene sets from distinct cell populations within the spatial expression profiles. The results revealed that cluster 0 was highly associated with fiber cells (**Supplementary Fig. 6**). Additionally, integration of the two datasets using Spotlight^36^ further demonstrated that genes specific to cluster 0 were predominantly localized to fiber cells in situ, while also providing spatial insights into the potential identities of other clusters (**Supplementary Fig. 7**). Notably, clusters 11 and 12, which were identified in the single-cell dataset, were not captured in the spatial transcriptome data due to the absence of bract tissue in the spatial samples, which comprised a smaller tissue region compared to the dissociated single-cell preparation. We next examined the spatial expression patterns of the top marker genes for each of the 13 cell clusters and observed that these genes exhibited highly specific expression in their corresponding cell types (**Supplementary Figs. 8-12**). Through these complementary approaches, we ultimately identified six distinct cell types: cluster 0 as fiber cells; clusters 1, 2, and 7 as ovary wall outer epidermal cells; cluster 3 as ovule cells; cluster 4 as placenta cells; clusters 5, 6, 8, and 9 as ovary wall parenchyma cells; and cluster 10 as nucellus cells (**Fig. 1B**). To further validate the accuracy of this cell type annotation, we performed RNA in situ hybridization for representative marker genes of each cell type (**Fig. 1D and 1E**). The results were consistent with the annotation, further confirming the reliability of the spatial transcriptome data. We further analyzed the functional pathways enriched in fiber cells (cluster 0). GO enrichment analysis revealed their involvement in ribosome, peptide biosynthetic process, and peptide secondary metabolic process (**Supplementary Fig. 13 and Fig. 14) (Supplementary Data 1**), consistent with the high demand for protein synthesis during fiber development and with features reported for cotton fiber initiation^16^.

### Transcriptional dynamics drive fiber cell functional transitions

Fiber cells undergo distinct morphological transitions during development, progressing through initiation, protrusion, and elongation stages. To investigate how dynamic gene expression underlies these functional changes, we further constructed single-cell transcriptomic maps for the initiation, protrusion, and elongation stages separately (**Fig. 2A-C**). UMAP dimensionality reduction analysis revealed a marked increase in the number of fiber cells at the protrusion and elongation stages compared to the initiation stage (**Fig. 2D**). Notably, although fiber cells were less abundant at the initiation stage, this population was already clearly defined at the transcriptional level. Consistent with this observation, *PtoMYB38* and *PtoSTK2*, two genes previously reported to regulate poplar fiber initiation^22,23^, were specifically and highly expressed at the initiation stage (**Fig. 2E**).

**Fig. 2.**
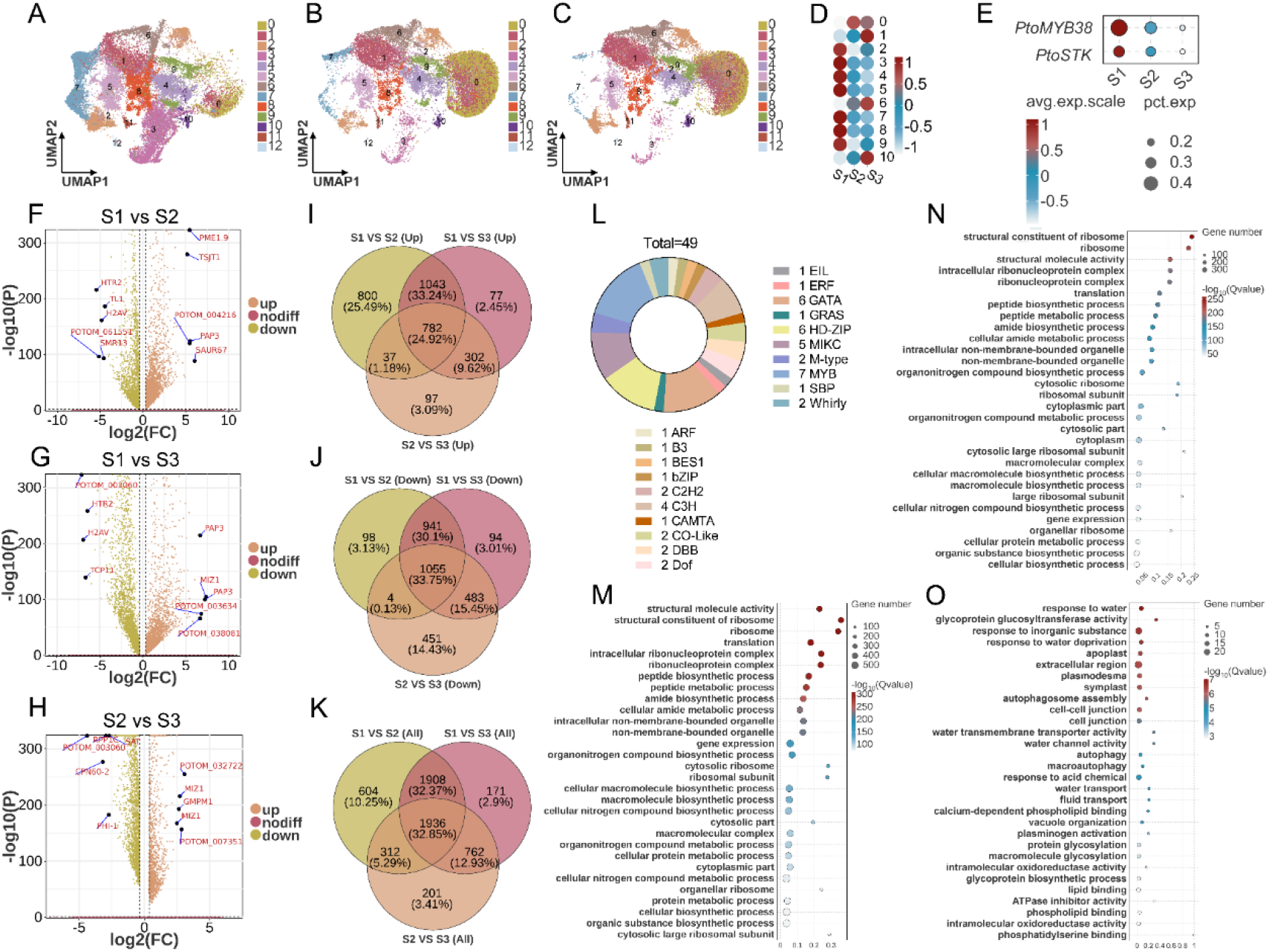
| Gene expression dynamics during fiber development. (**A**) Single-cell transcriptomic atlas of the capsule at stage 1 (initiation). (**B**) Single-cell transcriptomic atlas of the capsule at stage 2 (protrusion). (**C**) Single-cell transcriptomic atlas of the capsule at stage 3 (elongation). (**D**) Proportional changes of each cell cluster across developmental stages. (**E**) Expression patterns of *PtoMYB38* and *PtoSTK* across the three stages. (**F**) Differentially expressed genes in fiber cells at stage 2 relative to stage 1 (upregulated genes in orange, downregulated genes in green). (**G**) Differentially expressed genes in fiber cells at stage 3 relative to stage 1 (upregulated genes in orange, downregulated genes in green). (**H**) Differentially expressed genes in fiber cells at stage 3 relative to stage 2 (upregulated genes in orange, downregulated genes in green). (**I**) Venn diagram analysis to identify commonly upregulated DEGs in fiber cells across the three stages. (**J**) Venn diagram analysis to identify commonly downregulated DEGs in fiber cells across the three stages. (**K**) Venn diagram analysis to identify all commonly regulated DEGs in fiber cells across the three stages. (**L**) The 49 core transcription factors identified among the commonly regulated DEGs across all three stages. (**M**) GO enrichment analysis of stage-specific upregulated genes at stage 1. (**N**) GO enrichment analysis of stage-specific upregulated genes at stage 2. (**O**) GO enrichment analysis of stage-specific upregulated genes at stage 3.

To dissect the molecular dynamics underlying fiber cell development, we performed comparative analysis of differentially expressed genes (DEGs) across the three stages (**Fig. 2F–H**), identifying 782 commonly upregulated DEGs (accounting for 24.92% of all upregulated genes), 1,055 commonly downregulated DEGs (33.75% of all downregulated genes), and a total of 1,936 commonly regulated DEGs (32.85% of all DEGs) across all three stages (**Fig. 2I–K) (Supplementary Data 2-4**), indicating that a substantial proportion of genes are continuously involved throughout the entire process of fiber cell development from initiation through elongation. Given that transcription factors (TFs) play pivotal regulatory roles in plant development by modulating gene expression, we further investigated TF dynamics and identified a total of 49 core TFs differentially expressed across all three stages (**Supplementary Data 5**), belonging to families such as HD-ZIP, MYB, and GATA, suggesting their potential key regulatory functions in fiber cell development (**Fig. 2L**). To uncover functional transitions during fiber cell development, we first observed that the numbers of upregulated genes differed markedly among the three stages (**Supplementary Fig. 15**), prompting us to perform GO enrichment analysis on the stage-specific upregulated gene sets (**Supplementary Data 6**). We found that at the initiation stage, fiber cells were significantly enriched in pathways related to ribosome biogenesis and peptide biosynthetic process (**Fig. 2M**); at the protrusion stage, similar pathways were enriched, albeit with markedly altered gene participation ratios compared to the initiation stage (**Fig. 2N**); in contrast, at the elongation stage, pathways associated with environmental interactions, such as response to water and response to inorganic substances, became predominantly enriched (**Fig. 2O**). To further explore the global expression dynamics, we performed trend analysis on all genes expressed in fiber cells, which revealed eight distinct expression patterns. Notably, trend 0 (genes with highest expression at stage 1 followed by gradual decrease) and trend 7 (genes with lowest expression at stage 1 followed by gradual increase) exhibited opposite expression trajectories, accompanied by a corresponding functional shift in their enriched pathways (**Supplementary Fig. 16**). Collectively, these results demonstrate that temporal dynamics of gene expression drive the functional specialization and transitions of fiber cells as they progress through initiation, protrusion, and elongation.

### Developmental trajectory of fiber cells

To further elucidate the cellular origin of fiber cells and their functional transitions during development, we performed pseudotemporal analysis using Monocle 2 on fiber cells and placenta cells (**Fig. 3A**). The pseudotemporal trajectory revealed that both cell types reside along a common linear developmental path (**Fig. 3B**), with placenta cells distributed at the trajectory origin (**Fig. 3C**) and fiber cells positioned at the terminal end (**Fig. 3D**), confirming that fiber cells originate from placenta cells. This finding was further validated using Monocle 3, which also showed a gradual transition from placenta cells to fiber cells (**Fig. 3E–F**). To uncover the functional transitions underlying fiber cell development, we performed modular analysis of genes dynamically expressed along the pseudotemporal trajectory. The results revealed distinct functional modules associated with different developmental phases: genes in module C2, enriched at the trajectory origin, were significantly associated with developmental processes including unicellular organism development; as the trajectory progressed, genes in module C3 became enriched for molecular functions such as organic cyclic compound binding; at the trajectory terminus, genes in module C1 were predominantly enriched in pathways related to protein synthesis, including peptide biosynthetic process, peptide metabolic process, and ribosome (**Fig. 3G) (Supplementary Data 7**). Collectively, these results suggest that the differentiation from placenta cells to fiber cells may be orchestrated by dynamic changes in gene expression, potentially driving a continuous transition from developmental initiation toward functional specialization. To further explore the potential developmental trajectory of the capsule, we performed RNA velocity analysis on all cell types identified in the capsule using spatial transcriptome data. Multiple analytical approaches suggested the existence of a major developmental trajectory originating from ovary wall outer epidermal cells, with progressive transitions to ovary wall parenchyma cells and placenta cells, ultimately giving rise to ovule and fiber cells (**Supplementary Fig. 17 and Fig. 18**). This inference, although based solely on computational prediction, places the ovary wall outer epidermis near the root of the inferred capsule developmental hierarchy, raising the possibility that it may serve as the originating cell population for the capsule structure. However, these findings remain correlative and await experimental validation.

**Fig. 3.**
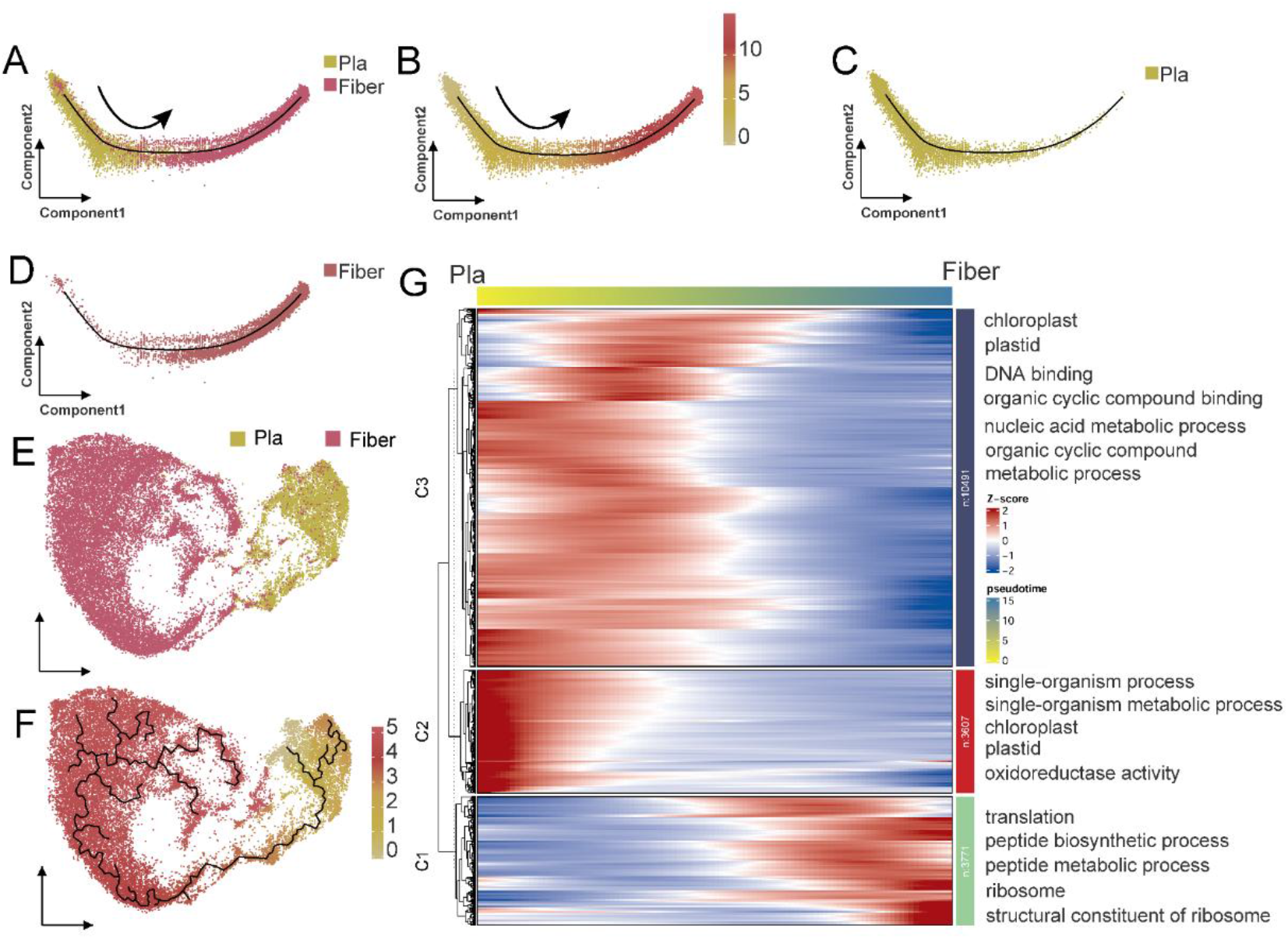
| Developmental trajectory of fiber cells. (**A**) Pseudotemporal trajectory analysis of placenta cells and fiber cells using Monocle 2. (**B**) Heatmap showing the pseudotemporal ordering of cells along the trajectory generated by Monocle 2, with the arrow indicating trajectory direction. (**C**) Distribution of placenta cells along the pseudotemporal trajectory, localized at the trajectory origin. (**D**) Distribution of fiber cells along the pseudotemporal trajectory, positioned at the trajectory terminus. (**E**) Pseudotemporal trajectory analysis of placenta cells and fiber cells using Monocle 3. (**F**) Heatmap displaying the pseudotemporal ordering of cells generated by Monocle 3. (**G**) Heatmap showing the expression dynamics of key genes along the fiber cell developmental trajectory. GO enrichment analysis of preferentially expressed genes along the trajectory is shown on the right, revealing their associated biological functions.

### Three subtypes of fiber cells

Different cell types, or even distinct subpopulations within a single cell type, can perform divergent functions, raising the possibility that fiber cells may also exhibit functional heterogeneity across developmental stages. To dissect the cellular heterogeneity and functional specialization within the fiber cell population, we performed unsupervised clustering analysis on all fiber cells and identified three transcriptionally distinct subtypes (subtypes 0, 1, and 2) (**Fig. 4A**). To reconstruct the transcriptional trajectory underlying fiber cell diversification, we performed pseudotemporal analysis using Monocle 2. Subtype 1 was predominantly enriched at stage 1 and localized to the left side of the trajectory (**Fig. 4B**); therefore, we designated this region as the trajectory origin (**Fig. 4C**). This developmental path was further validated using Monocle 3 (**Fig. 4D and 4E**), and CytoTRACE analysis confirmed that subtype 1 exhibits the highest differentiation potential (**Fig. 4F**). Collectively, these results establish a developmental hierarchy in which fiber cells originate from subtype 1 and progressively differentiate into subtypes 0 and 2.

**Fig. 4.**
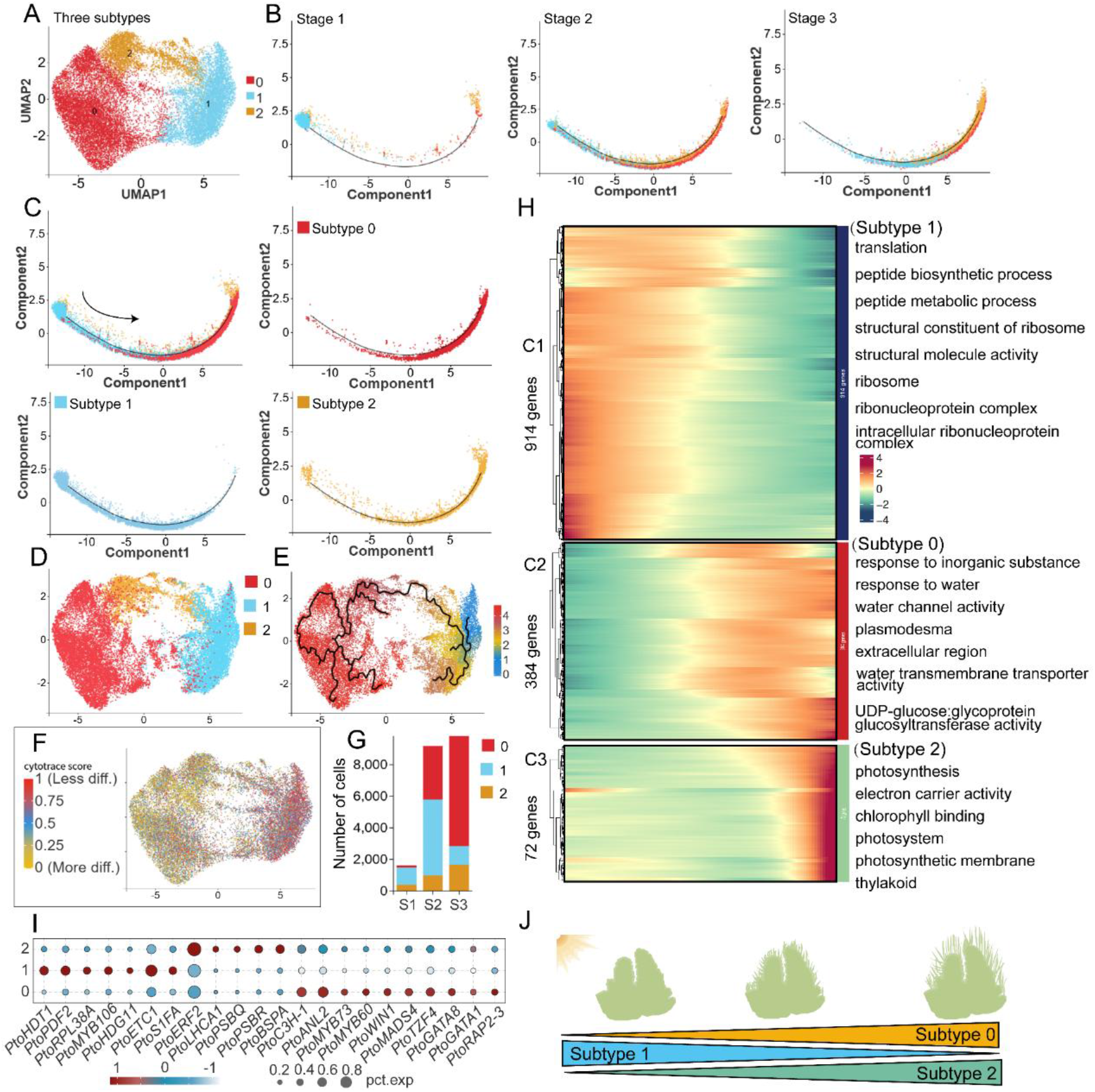
| Reclustering and pseudotemporal trajectory of fiber cells. (**A**) Reclustering of fiber cells identifies three putative subtypes (subtypes 0, 1, and 2). (**B**) Pseudotemporal trajectories of fiber cells at the initiation (stage 1), protrusion (stage 2), and elongation (stage 3) stages. (**C**) Pseudotemporal trajectory analysis of the three fiber cell subtypes (subtypes 0, 1, and 2) inferred by Monocle 2, showing both the combined trajectory of all three subtypes and the individual trajectories of each subtype. (**D**) Visualization of the three fiber cell subtypes using Monocle 3. (**E**) Pseudotemporal trajectory analysis of the three fiber cell subtypes using Monocle 3. (**F**) CytoTRACE analysis showing the differentiation potential of the three fiber cell subtypes. (**G**) Number of cells in each fiber cell subtype across the three developmental stages. (**H**) Heatmap displaying the expression patterns of core genes across the three fiber cell subtypes. GO enrichment analysis of preferentially expressed genes in each subtype is shown on the right, revealing their associated biological functions. (**I**) Bubble plot showing specifically highly expressed genes in each of the three fiber cell subtypes. (**J**) Schematic model illustrating the temporal coordination of the three fiber cell subtypes during fiber development.

Interestingly, all three subtypes were present at each developmental stage, albeit with markedly dynamic shifts in their relative proportions (**Fig. 4B and 4G**). Subtype 1 showed a sharp increase from the initiation to protrusion stage, a critical window for fiber cell fate determination and protrusion onset, suggesting its primary role in initiating fiber cell specification. In contrast, subtypes 0 and 2 exhibited gradually increasing proportions as development progressed, implicating their involvement in later phases of fiber development (**Fig. 4C**). To uncover the functional specialization of each subtype, we analyzed their differentially expressed genes and enriched pathways (**Fig. 4H) (Supplementary Data 8**). Subtype 1 was specifically enriched in pathways related to peptide biosynthetic process and ribonucleoprotein complex, a transcriptional signature consistent with cotton fiber initials^16^, further supporting its role as the initiating subtype. Subtype 0 showed enrichment in pathways associated with response to inorganic substances, response to water, and plasmodesmata, processes linked to cell elongation and homeostasis regulation, suggesting its primary involvement in fiber elongation. Subtype 2 was significantly enriched in photosynthesis-related pathways, including photosynthesis, chlorophyll binding, and thylakoid, indicating a role in energy provision and environmental signaling during fiber development. Analysis of candidate regulatory genes further supported this functional division (**Fig. 4I) (Supplementary Data 9**). Subtype 1 highly expressed transcription factors from the HD-ZIP family (including *PtoPDF2* and *PtoHDG11*), as well as *PtoHDT1*, *PtoETC1*, *PtoS1FA*, *PtoRPL38A*, and MYB family members such as *PtoMYB106*, with *PtoMYB106* previously implicated in poplar fiber initiation^23^. Subtype 2 exhibited high expression of a set of photosynthetic genes, including LHC family members such as *PtoLHCA1*, as well as multiple genes belonging to the PsaA and PsaB families, including *PtoPSBQ*, *PtoPSBR*, *PtoBSPA*, and *PtoPSAL*, and the transcription factor *PtoERF2*, corroborating its photosynthetic function. Subtype 0 was enriched for multiple transcription factors, including *PtoANL2*, *PtoWIN1*, and *PtoMADS4*, suggesting its functions involve coordinated regulation by multiple factors.

Based on these findings, we propose a working model of cooperative fiber cell subtype dynamics (**Fig. 4J**): Subtype 1 acts as the initiating population, responsible for fiber cell fate determination; subsequently, subtype 2 becomes activated, providing energy through photosynthesis and sensing environmental cues; finally, subtype 0 takes over, driving rapid fiber elongation. The temporal coordination and dynamic shifts in subtype proportions collectively orchestrate the complete developmental program of fiber cells, offering new insights into how cellular heterogeneity contributes to plant cell differentiation.

### Identification of key regulators governing fiber cell development

To elucidate cell type-specific regulatory networks underlying fiber cell development in poplar capsules, we performed WGCNA and identified 21 distinct modules (**Supplementary Fig. 19**). Correlation analysis revealed that three modules (turquoise, darkgreen, and darkgrey) were significantly associated with fiber cell development (**Supplementary Data 10-11**). Specifically, the turquoise module showed high correlation with fiber cell initiation at stage 1, whereas the darkgreen and darkgrey modules were more closely associated with fiber elongation at stages 2 and 3.

KEGG enrichment analysis of the initiation-associated turquoise module revealed significant enrichment in ribosome, proteasome, and protein export pathways (**Supplementary Fig. 20**), consistent with the notion that fiber cells, upon differentiation from placenta cells, require extensive protein synthesis machinery to support subsequent protrusion and elongation. This aligns with the molecular features of cell fate determination in cotton fiber initials, where downregulation of cell cycle genes is coupled with upregulation of ribosome biosynthesis and translation-related genes^37^. Gene network analysis further identified *PtoPDF2*, *PtoMYB38*, and *PtoRPL38A* as hub genes within this module (**Supplementary Fig. 19**). Among these, *PtoMYB38* has been previously characterized as a positive regulator of fiber initiation, as its loss-of-function leads to fiberless capsules^23^.

For the elongation-associated modules, KEGG enrichment analysis revealed that both darkgreen and darkgrey were enriched in fatty acid-related metabolic pathways, with darkgreen showing enrichment in fatty acid metabolism and metabolic pathways, and darkgrey specifically enriched in biosynthesis of unsaturated fatty acids (**Supplementary Fig. 20**). This enrichment pattern aligns with the established role of lipid metabolism in fiber elongation, as demonstrated in cotton where very-long-chain fatty acid (VLCFA) synthesis is essential for this process^38^. Collectively, these results demonstrate that the spatiotemporal co-expression regulatory network constructed from snRNA-seq data effectively captures key regulators of fiber cell development and provides candidate targets for functional validation.

### PtoPDF2 targets known seed fiber regulators *PtoMYB38* and *PtoSTK2* in poplar

Through WGCNA of the single-nucleus transcriptome data, we identified the turquoise module as highly correlated with fiber cell initiation (stage 1). Within this module, the HD-ZIP IV transcription factor gene *PtoPDF2* emerged as a hub gene with high intramodular connectivity (**Supplementary Fig. 19**). Consistently, our earlier subtyping analysis of fiber cells revealed that *PtoPDF2* is preferentially expressed in subtype 1, the subpopulation that marks the initiation stage and is responsible for cell fate determination (**Fig. 4N**). These complementary lines of evidence position PtoPDF2 as a strong candidate regulator of early fiber cell development. To understand the evolutionary context of PtoPDF2, we first constructed a phylogenetic tree of HD-Zip IV transcription factors from *Arabidopsis* and poplar. PtoPDF2 clustered most closely with AtPDF2 (Fig. 5A), indicating a close evolutionary relationship. To further explore its potential regulatory network, we examined the expression patterns of *PtoPDF2* together with two known fiber-related genes, *PtoMYB38* and *PtoSTK2*^22,23^. Across the 13 cell clusters, *PtoPDF2* and *PtoMYB38* were preferentially expressed in fiber cells (cluster 0), whereas *PtoSTK2* showed the highest expression in placenta cells (cluster 4) and moderate expression in fiber cells (cluster 0) (**Fig. 5B**). Notably, all three genes exhibited peak transcript levels at the initiation stage (stage 1) within fiber cells, with gradual decline during protrusion (stage 2) and elongation (stage 3) (**Fig. 5C**). This coordinated temporal pattern suggests that the three genes may participate in the same developmental process and potentially be governed by a common regulatory network during early fiber cell fate specification. Although the turquoise module also contained several GA signaling genes, LCA and BiFC assays detected no in vivo interaction between PtoPDF2 and the DELLA repressor GIBBERELLIC ACID INSENSITIVE (PtoGAI) (**Supplementary Fig. 21**), suggesting that PtoPDF2 may act independently of this core GA repressor under our experimental conditions.

**Fig. 5.**
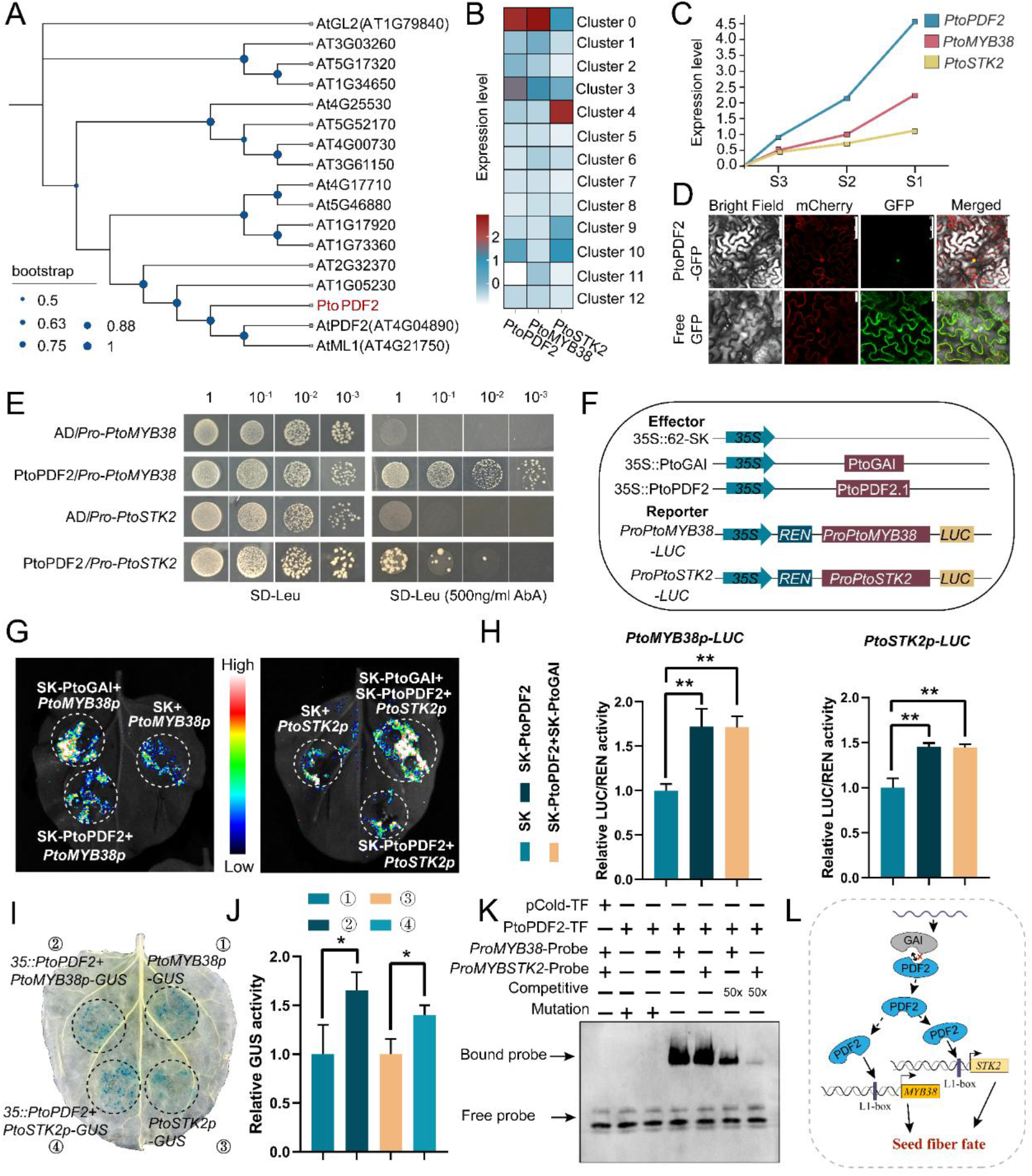
| PtoPDF2 acts as an upstream regulator of *PtoMYB38* and *PtoSTK2* during poplar seed fiber development. (**A**) Phylogenetic analysis of HD-Zip IV transcription factors from *Arabidopsis* and poplar. PtoPDF2 clusters most closely with AtPDF2. (**B**) Expression heatmap of *PtoPDF2*, *PtoMYB38* and *PtoSTK2* across the 13 distinct cell clusters identified in poplar capsules. (**C**) Expression dynamics of *PtoPDF2*, *PtoMYB38* and *PtoSTK2* within fiber cells (cluster 0) at the initiation (stage 1), protrusion (stage 2) and elongation (stage 3) stages. (**D**) Subcellular localization of PtoPDF2 in *N. benthamiana* epidermal cells. The PtoPDF2-EGFP fusion protein localizes exclusively to the nucleus, whereas the free EGFP control shows diffuse cytoplasmic and nuclear signal. Scale bars, 75 μm (upper) and 25 μm (lower). (**E**) Yeast one-hybrid (Y1H) assays demonstrating that PtoPDF2 binds to the promoter fragments of *PtoMYB38* and *PtoSTK2*. Transformants grown on SD/-Leu medium containing 500 ng mL⁻¹ Aureobasidin A (AbA) indicate positive interactions. (**F**) Schematic representation of the effector and reporter constructs used in dual-luciferase assays. Effectors: PtoPDF2 and PtoGAI driven by the CaMV 35S promoter. Reporters: firefly luciferase driven by the *PtoMYB38* or *PtoSTK2* promoter. (**G-H**) Dual-luciferase reporter assays showing that PtoPDF2 activates the promoters of *PtoMYB38* and *PtoSTK2*. Co-expression of PtoGAI with PtoPDF2 does not alter this binding. (**I-J**) GUS reporter assays showing that PtoPDF2 binds to the promoters of *PtoMYB38* and *PtoSTK2* and promotes their expression. (**K**) Electrophoretic mobility shift assays (EMSA) confirming that PtoPDF2 directly binds to the L1-box motifs in the *PtoMYB38* and *PtoSTK2* promoters. (**L**) Proposed working model: PtoPDF2 binds to the L1-box elements in the promoters of *PtoMYB38* and *PtoSTK2*, thereby regulating seed fiber development. Data are mean ± S.D. of three independent biological replicates, two-tailed Student’s *t*-test. \*\**P* < 0.01, \**P* < 0.05.

Subcellular localization confirmed that PtoPDF2-EGFP was exclusively localized to the nucleus (**Fig. 5D**), consistent with its role as a transcription factor. To test whether PtoPDF2 directly acts on the promoters of *PtoMYB38* and *PtoSTK2*, we first performed yeast one-hybrid (Y1H) assays. Transformants carrying PtoPDF2 grew on selective medium containing 500 ng mL⁻¹ Aureobasidin A (**Fig. 5E**), confirming that PtoPDF2 specifically binds to both target gene promoters. To further assess the transcriptional activation activity, we performed dual-luciferase reporter assays. Effector constructs expressed PtoPDF2 under the CaMV 35S promoter, while reporter constructs carried firefly luciferase driven by the *PtoMYB38* or *PtoSTK2* promoter (**Fig. 5F**). The results showed that PtoPDF2 activates the expression of both reporters, while co-expression of PtoGAI did not alter this activation (**Fig. 5G-H**). These findings were further corroborated by GUS reporter assays, which produced consistent results (**Fig. 5I-J**). To pinpoint the cis-elements mediating this regulation, we performed electrophoretic mobility shift assays (EMSA), which confirmed that PtoPDF2 directly binds to the *L1-box* motifs in both promoters (**Fig. 5K).** Based on these findings, we propose a working model in which PtoPDF2 binds to L1-box elements in the promoters of *PtoMYB38* and *PtoSTK2*, thereby contributing to the regulatory network that governs seed fiber development (**Fig. 5L**).

## Discussion

Seed fibers are the defining cell type of the female reproductive organ in *Populus* and the direct cause of the annual “fluff” nuisance. Yet they remain one of the few major plant cell types with no defined cellular origin and no regulatory framework. Work in this species has advanced mainly by testing homologs of genes first characterized in *Arabidopsis* or cotton^24,39^ and by bulk transcriptomics^25^. These approaches have provided valuable gene resources and insights into conserved regulatory mechanisms. However, they average across the entire capsule. They therefore cannot separate the transcriptional events in the few hundred epidermal cells that will become fibers from those in the surrounding wall, ovule, and nucellus. Single-cell and spatial technologies have begun to resolve cell-type heterogeneity in vegetative organs of *Arabidopsis*^40^ and rice^41^ and in the roots^42^, stems^43^, and leaves^44^ of *Populus* itself. Recently, the first spatiotemporal atlases of reproductive development have also appeared in conifers^45^ and in the cotton fiber system^33^. What has been missing is the equivalent for seed fiber development in a woody perennial: a dataset in which the cells that will become fibers are resolved as a population, placed in their tissue context, and followed across the morphological transitions that define their differentiation. Here we address this gap by resolving 13 cell clusters from more than 80,000 nuclei across the three stages of fiber morphogenesis and anchoring them to the capsule structure with spatial transcriptomics (**Fig. 1B**). To our knowledge, this is the first spatiotemporal single-cell atlas of trichome development on reproductive organs in any woody perennial, and the first in which the regulatory logic of fiber initiation is derived from the fiber cells themselves rather than inferred from other species. Fluff is a reproductive trait, and efforts to manipulate it have been limited by the lack of a cell-type-resolved target. The molecular module we identify during fiber development (see below) therefore provides a concrete entry point for breeding low-fluff cultivars. More broadly, it shows that single-cell resolution is not a technical refinement here, but a prerequisite for asking the right question.

The most important finding of this study concerns the accurate origin of seed fibers in poplar. Cotton is the reference system for single-celled seed fibers. Its lint and fuzz arise from the epidermis of the ovule outer integument, a maternal seed coat layer, and the entire genetic framework of fiber initiation has been built on that anatomical premise^46,47^. Poplar fibers arise in a different place: the adaxial epidermis of the placenta, the ridge from which the ovules themselves are initiated. Three independent lines of evidence support this conclusion: the position of new fibers in histological sections (**Supplementary Fig. 1**), the pseudotime trajectories computed from placenta and fiber cells (**Fig. 3**), and the RNA velocity field reconstructed across the capsule (**Supplementary Fig. 18**). The placenta epidermis is not simply a supporting layer. It is the same cell layer in which a placenta-limited *MIXTA* gene drives epidermal differentiation through an MBW-type complex^24^. The fiber program is therefore built on an epidermis that is already able to adopt a trichome-like fate. This finding resolves a long-standing question in poplar reproductive biology and changes how seed fibers should be compared across species. Cotton and poplar fibers are not homologous cell populations. They were recruited independently from two different maternal epidermal layers of the reproductive organ. This distinction matters in practice. Regulatory hypotheses imported from cotton, for example that ovule-expressed *MIXTA*-like or *MYB* genes act cell-autonomously in fiber initials, cannot be assumed to transfer. In poplar, the factor responsible must therefore be present in the placenta epidermis, or must send a signal to it, before ovule formation is complete. Our model also implies that the early steps of the poplar fiber program may be shared with placenta development rather than with the ovule outer integument. This can be tested by comparing placenta epidermal cells between fiber-bearing and fiberless genotypes.

A second unexpected result is that the fiber population is not homogeneous. Single-cell clustering resolves three transcriptionally distinct subtypes. All three are present at every stage, but their proportions shift as development proceeds. The initiating subtype is enriched for *HD-ZIP* and *MYB* regulators and for peptide biosynthetic and ribonucleoprotein machinery. The elongation subtype is enriched for water and inorganic-substance responses and for plasmodesmatal connectivity. The third subtype has a photosynthetic signature, comprising *LHC*, *Psa* and *Psb* genes together with *PtoERF2* (**Fig. 4**). In cotton, a division of labor between fiber initials and elongating fibers is well established: fate determination precedes polar elongation4^47–49^, and fatty acid and cell-wall programs sustain rapid expansion^50,51^. A photosynthetic subtype within a seed fiber population has not been described before, to our knowledge. Its presence is unexpected, because the fiber cells differentiate inside a closed capsule whose wall is still green, and because a rapidly elongating cell is usually assumed to depend on the maternal plant for its energy. This subtype may therefore contribute to the energy needed for elongation, or it may sense conditions inside the capsule and help time fiber expansion to capsule maturation. Both possibilities remain to be tested. That all three subtypes are present from the earliest stage sampled argues that they are not simply sequential states of one cell, but a division of labor established at or very close to fate specification. One morphologically defined cell type can therefore contain several transcriptionally and functionally distinct subpopulations, as shown in root columella, root epidermis and leaf^31,52–54^. Seed fibers extend this principle to the reproductive organ of a woody perennial. Whether the subtypes are fixed or can interconvert, a question that bears directly on breeding strategies aiming to reduce fiber number rather than block elongation, will require lineage tracing in planta.

We summarize our findings as a working model of fiber cell subtype dynamics across the three developmental stages (**Fig. 6**). In this model, the fiber population is not a uniform cell type that matures as a whole, but a shifting mixture of three transcriptionally distinct subtypes, with each stage dominated by a different population and transcriptional program. The initiating subtype is defined by the PtoPDF2–L1-box– *PtoMYB38*/*PtoSTK2* module together with *PtoRPL38A* and the SPL-family gene *PtoSPL6*, and is most prominent at initiation. The photosynthetic subtype expands most rapidly during protrusion. The elongating subtype, marked by *PtoANL2*, *PtoWIN1*, *PtoMADS4*, *PtoGATA8* and *PtoMYB60*, accumulates during elongation as fibers come to occupy most of the capsule interior. The core of the model is a simple cascade. The HD-Zip IV transcription factor PtoPDF2 binds L1-box elements^55^in the promoters of *PtoMYB38* and *PtoSTK2* and activates their expression. It therefore acts upstream of two genes whose loss of function abolishes fiber formation^22,23^. The L1-box and its recognition by HD-Zip IV proteins are conserved^56,57^, but this module has not been described in a reproductive organ of a woody plant before. The genes shown for each stage in Fig. 6 are the next layer of this network. They are candidate regulators, proposed on the basis of stage-restricted expression and co-expression rather than demonstrated causality, and they remain to be tested.

**Fig. 6.**
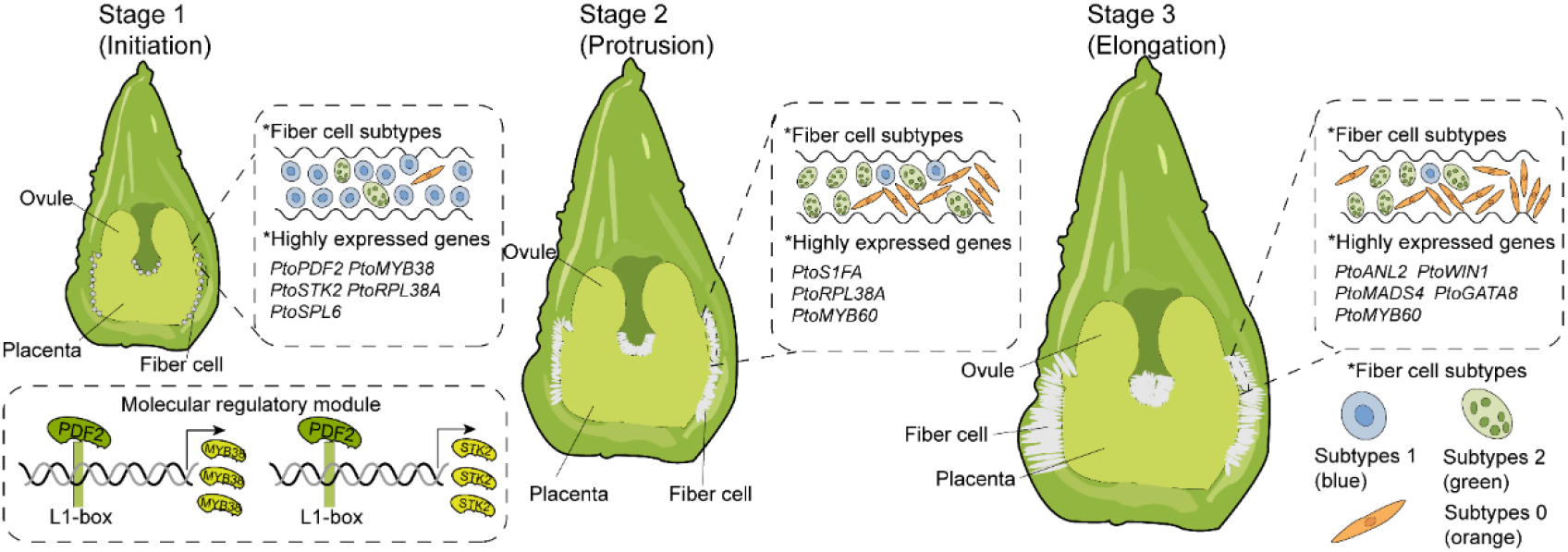
| A working model of fiber cell subtype dynamics and the PtoPDF2–L1-box regulatory module in the developing poplar capsule. Fiber cell development in the female capsule proceeds through initiation, protrusion and elongation (Stages 1–3). The fiber population comprises three transcriptionally distinct subtypes (key at bottom right): subtype 1 (blue) predominates at initiation and is progressively supplemented by subtype 2 (green), which expands most rapidly during protrusion, and by subtype 0 (orange), which becomes the most abundant population during elongation. Representative stage-restricted genes are indicated for each stage. Inset, PtoPDF2 binds L1-box elements in the promoters of *PtoMYB38* and *PtoSTK2* and activates their transcription, defining the regulatory core of fiber development.

In a comparative context, this module shows how differently plants have solved the problem of epidermal specialization. *Arabidopsis* leaf trichome fate is governed by an R2R3 MYB–bHLH–WD40 (MBW) complex that is modulated by R3 MYB competitors and phytohormones^9,39,58,59^. Tomato and cucumber instead use interacting *HD-Zip IV* and *C2H2* zinc-finger regulators ^60–63^, and cotton fiber initiation is controlled by *MIXTA*-like and *MYB25*-like factors^64–68^. Poplar seed fiber occupies an intermediate position: it uses MBW-type effectors as terminal determinants, but places an HD-Zip IV factor upstream of them. These two layers can therefore be wired in series rather than representing alternative strategies. *PtoPDF2* is most closely related to *AtPDF2* (**Fig. 5A**), and HD-Zip IV proteins are not all pro-trichome in poplar: *PtoHDG11* is co-expressed in initiating fibers, yet its overexpression suppresses trichome formation^69^. The evolutionary novelty therefore appears to lie less in the regulators themselves, most of which have recognizable homologs elsewhere, than in how they are connected and which targets they control. Single-cell atlases of vegetative and reproductive organs are now available across species^33,40^. Comparative single-cell analyses may in future show whether this arrangement reflects an ancient module that has been reused many times for epidermal specialization, or a convergence specific to the reproductive organs of woody plants.

This study provides a single-cell, spatially resolved framework for seed fiber development in a woody perennial and defines a module (PtoPDF2 – L1-box –*PtoMYB38*/*PtoSTK2*) that acts in the initiating fiber subpopulation (**Fig. 6**). Seed fibers arise from the placenta epidermis rather than the ovule integument, the fiber population comprises at least three cooperating subtypes, and fiber development is controlled by a compact and experimentally accessible regulatory module. Together these results turn an environmentally important trait from a phenological problem into a defined molecular target.

## Methods

### Plant materials and growth conditions

Branches of female *P*. *tomentosa* were collected from the nursery of Beijing Forestry University, Beijing, China (39.9 ° N, 116.3 ° E), in late February. At the time of collection, flower buds had completed the natural chilling of the Beijing winter and had been released from endodormancy; branches therefore flowered within days of being transferred to the growth chamber, and no additional chilling treatment was applied. The branches were subsequently hydroponically cultured in a controlled-environment growth chamber set to 25°C with a photoperiod of 16 h light/8 h dark. Based on our previous characterization of poplar capsule developmental stages^4^, capsules were sampled from female inflorescences at 8-hour intervals from 3 to 6 days post-anthesis (DPA). At each time point, capsules were either embedded in optimal cutting temperature (OCT) compound or flash-frozen in liquid nitrogen for subsequent analyses. For *in situ* hybridization and paraffin sectioning, capsules were immediately fixed in *in situ* hybridization fixative (formalin, glacial acetic acid, and 70% ethanol, treated with DEPC to inactivate RNases) and FAA fixative (5% formalin, 5% glacial acetic acid, and 90% of 50% ethanol), respectively.

### Cytological observation by paraffin sectioning

FAA-fixed capsule samples were dehydrated through a graded ethanol series (50%, 70%, 80%, 90%, 95%, and 100%, v/v), followed by clearing with a graded xylene series (25%, 50%, 75%, and 100% xylene in ethanol, v/v). The samples were then infiltrated with paraffin at 60°C for three days, with the paraffin solution replaced three times during this period. Subsequently, the samples were embedded in paraffin blocks. Serial sections of approximately 10 μm in thickness were obtained using a rotary microtome (LEICA RM2265, Leica, Germany). The sections were dewaxed in xylene, rehydrated through a graded ethanol series, and stained with safranin O and Fast Green FCF. Finally, the stained sections were observed and imaged under a light microscope (Echo Revolve, ECHO, San Diego, CA, USA).

### RNA in situ hybridization

RNA in situ hybridization was performed to validate the spatial expression patterns of selected marker genes. Capsule samples were fixed in DEPC-treated formalin-acetic acid-70% ethanol, dehydrated, and embedded in paraffin. Sections of 8 μm thickness were prepared for hybridization. Digoxigenin (DIG)-labeled antisense and sense probes were synthesized using a DIG RNA labeling kit (Roche, Switzerland) according to the manufacturer’s instructions. The primer sequences used for probe synthesis are listed in Supplementary Table 4. Hybridization was performed overnight at 55°C in hybridization buffer containing 50% formamide. After stringent washes, immunological detection was carried out using alkaline phosphatase-conjugated anti-DIG antibody (Roche, Switzerland). Each gene was detected in at least three independent biological replicates.

### Single-nucleus isolation and snRNA-seq library preparation

Frozen capsule samples collected at three key developmental stages (4 DPA 24:00, 5 DPA 24:00, and 6 DPA 16:00) with three biological replicates per stage were used for single-nucleus RNA sequencing. Nuclei isolation was performed following established protocols with modifications. Briefly, frozen tissues were finely chopped with a sterile razor blade for 2 min, repeated 2-3 times until homogenized, and transferred to pre-chilled Nuclei Isolation Buffer (NIB) containing 5% Dextran T40, 0.4 M sucrose, 10 mM MgCl₂, 1 mM DTT, 2 U/μL RNase inhibitor, 0.1% Triton X-100, and 100 mM Tris-HCl (pH 7.4). The homogenate was centrifuged at 300 × g for 1 min at 4°C, and the supernatant was collected. The supernatant was sequentially filtered through 70 μm and 40 μm cell strainers, followed by centrifugation at 2000 × g for 5 min at 4°C. The pellet was resuspended in Wash Buffer (1× PBS, 1% BSA, and 2 U/μL RNase inhibitor).

For fluorescence-activated nuclei sorting (FANS), the nuclear suspension was stained with DAPI at a final concentration of 10 μM. Sorting was performed on a flow cytometer (BD FACSAria III, BD Biosciences, USA) equipped with a 70 μm nozzle at 20 psi. Nuclei were gated based on FSC-A/SSC-A to exclude debris, and DAPI-positive nuclei were sorted into collection tubes containing Wash Buffer. Sorted nuclei were centrifuged at 2000 × g for 5 min at 4°C, resuspended in Wash Buffer, and counted using a hemocytometer under a fluorescence microscope. The final nuclear concentration was adjusted to approximately 1000 nuclei/μL.

Purified nuclei were immediately loaded onto the 10x Genomics Chromium Controller for droplet generation. snRNA-seq libraries were constructed using the Chromium Next GEM Single Cell 3′ Reagent Kits v3.1 according to the manufacturer’s instructions. Library quality was assessed on an Agilent 2100 Bioanalyzer using High Sensitivity DNA chips, and sequencing was performed on an Illumina NovaSeq 6000 platform with a paired-end 150 bp read length.

### Single-nucleus RNA-seq data processing and quality control

Raw sequencing data were demultiplexed and converted to FASTQ format using the bcl2fastq software (Illumina). Sample demultiplexing, barcode processing, and UMI counting were performed using the Cell Ranger Single Cell Software Suite (v6.1, 10x Genomics)^70^. Reads were aligned to the *P. tomentosa* reference genome ^71^ using the STAR aligner integrated into Cell Ranger. Only confidently mapped reads with valid barcodes and UMIs were retained for downstream analysis. The gene-barcode matrix was generated using the cellranger count pipeline. Initial quality control was performed to filter out low-quality cells. Cells with fewer than 200 detected genes or mitochondrial gene content exceeding 30% were excluded. Additionally, genes expressed in fewer than three cells were removed. Further filtering and normalization were conducted using the Seurat package (v4.0.4) in R. After quality control ^72^, the median genes per cell and median UMI counts per cell were calculated to assess data quality, confirming the suitability of the dataset for subsequent analyses.

### Spatial transcriptome profiling

Fresh capsule samples collected at stage 2 (5 DPA 24:00) were immediately embedded in Optimal Cutting Temperature compound (OCT, Sakura Tissue-TEK) on dry ice and stored at −80°C until sectioning. Tissue blocks were sectioned at a thickness of 10 μm using a pre-cooled cryostat, and sections were carefully transferred onto the 6.8 × 6.8 mm² oligo-barcoded capture areas of BMKMANU S3000 Gene Expression Slides (BMKMANU, ST03009). Each capture area contains approximately 4,140,000 spatially barcoded spots with a diameter of 2.5 μm and a center-to-center distance of 3.5 μm, providing an average of 7 to 37 spots per cell.

Prior to full-scale experiments, optimal tissue permeabilization conditions were determined using the BMKMANU S3000 Tissue Optimization Kit (ST03003) according to the manufacturer’s instructions. The fluorescent footprint was imaged using a Pannoramic MIDI slide scanning platform to evaluate permeabilization efficiency and select the optimal treatment duration. For gene expression analysis, tissue sections on the slides were fixed and stained with toluidine blue O (TBO) by incubating in isopropanol for one minute. Brightfield images were acquired at 40× magnification using a Pannoramic MIDI microscope to capture histological details of the tissue sections. Following imaging, tissue permeabilization was performed under the optimized conditions, and cDNA synthesis was carried out directly on the slide to capture spatially barcoded mRNA molecules.

Spatial transcriptome libraries were constructed using the BMKMANU Gene Expression Library Construction Kit (ST03002-34) following the manufacturer’s protocol. Library quality was assessed on an Agilent 2100 Bioanalyzer, and sequencing was performed on an Illumina NovaSeq X Plus platform with a paired-end 150 bp read length, achieving a minimum sequencing depth of 150,000 read pairs per spatial spot.

### Spatial transcriptome data processing

Raw sequencing data in FASTQ format were processed using BSTMatrix v2.0 (BMKMANU) for demultiplexing, barcode assignment, and generation of spatial gene expression matrices. The processed reads were aligned to the *Populus* reference genome using STAR aligner version 2.5.1b. For visualization of spatial gene expression, the TBO-stained histological images were manually aligned with the spatial transcriptome coordinates.

Downstream analysis was performed using the Seurat v4.0.1 package in R. Spatial spots with fewer than 300 detected genes or more than 30% mitochondrial gene content were filtered out as low-quality spots. Genes detected in fewer than five spatial spots were excluded from further analysis. Normalization was performed using the SCTransform function with the “assay = spatial” parameter to account for technical variation while preserving biological heterogeneity.

For cell segmentation and identification of spatial cell types, we performed multiscale cell segmentation using BSTMatrix v2.3j integrated with Cellpose v2.2.3. The TBO-stained images, which clearly visualized cell wall boundaries, were used as input for segmentation to define individual cell boundaries and assign spatial transcriptome spots to corresponding cells. Clustering analysis was performed at multiple resolution levels (L3, L6, L9, and L18), and level 6 (L6) was selected as the optimal resolution for downstream analysis based on its accurate representation of cellular heterogeneity in poplar capsules. Cell types were annotated by integrating spatial transcriptome data with snRNA-seq reference datasets using multimodal interaction analysis and the Spotlight algorithm^36^. The spatial distribution of specific cell types and marker genes was visualized using Seurat’s spatial feature plot functions^72,73^.

### Cell clustering and dimensionality reduction

To mitigate technical noise from individual genes in the single-nucleus transcriptomic data, dimensionality reduction was performed using principal component analysis (PCA) implemented in the Seurat package^74^. Principal components (PCs) representing coordinated gene expression patterns were selected for downstream analysis. To determine the optimal number of PCs, a resampling-based test was applied to evaluate the contribution of each PC to the overall dataset. PCs showing significant enrichment of low P-value genes were retained for subsequent clustering. Graph-based clustering was performed using Seurat’s shared-nearest neighbor (SNN) algorithm. Briefly, Euclidean distances between cells were calculated based on the selected PCs, and cells were embedded into an SNN graph. The graph was then partitioned into highly interconnected quasi-communities based on the Jaccard distance between neighboring cells, and cell clusters were identified using the Louvain algorithm for modularity optimization^75^. For visualization of cellular heterogeneity, nonlinear dimensionality reduction was applied. Both t-distributed stochastic neighbor embedding (t-SNE)^76^ and uniform manifold approximation and projection (UMAP)^77^were used to project high-dimensional single-cell data into two-dimensional space, where cells with similar transcriptional profiles cluster together while distinct populations are spatially separated, facilitating intuitive visualization of cellular diversity.

### Integration of single-nucleus and spatial transcriptomics data

To spatially map cell types identified from snRNA-seq data, we employed two complementary approaches. First, using the Seurat package (v4.0.1), we performed label transfer analysis^72,73^. The snRNA-seq dataset with defined cell type annotations was set as the reference. Anchor pairs between the reference and spatial transcriptomics data were identified using the FindTransferAnchors function with default parameters. Cell type probabilities for each spatial spot were then predicted using the TransferData function, and the cell type with the highest prediction score was assigned to each spot. Second, we applied the SPOTlight algorithm (v0.1.7) for spatial deconvolution^36^. Marker genes for each snRNA-seq cell cluster were first identified using the FindAllMarkers function in Seurat with parameters only.pos = TRUE, and genes with adjusted P-value < 0.1 were retained. These marker genes were then used as input for SPOTlight to deconvolute the spatial transcriptomic spots, generating a cell type proportion matrix for each spot. The cell type with the maximum proportion was assigned as the final identity for each spatial spot. Default parameters were used for all SPOTlight analyses. The spatial distribution of specific cell clusters and marker genes was visualized using Seurat’s spatial feature plot functions.

### Differential expression analysis and functional enrichment

For differential expression analysis between different samples or conditions within the same cell cluster, we first constructed a combined metadata label for each cell by concatenating its sample group information and cell cluster identity. Differential expression analysis was then performed using the Model-based Analysis of Single-cell Transcriptomics (MAST) framework^78^ implemented in the Seurat package^73^. MAST employs a hurdle model to account for both the fraction of cells expressing a gene and the expression level among expressing cells. Significance was assessed after multiple testing correction using the Benjamini-Hochberg method. Significantly differentially expressed genes were identified based on the following criteria: (1) absolute log2 fold change (|log₂FC|) ≥ 1; (2) adjusted P-value ≤ 0.05; and (3) minimum percentage of cells expressing the gene in the target population ≥ 10%. For identification of upregulated genes specific to individual subclusters, we used the FindAllMarkers function in Seurat with the following threshold parameters: minimum percentage of cells expressing the gene in either population (min.pct) > 0.25, P-value < 0.01, and fold change > 1.28.

To investigate the functional implications of differentially expressed genes, we performed Gene Ontology (GO)^79^ and Kyoto Encyclopedia of Genes and Genomes (KEGG)^80^ pathway enrichment analyses. For GO analysis, differentially expressed protein-coding genes were mapped to GO terms from the Gene Ontology database (http://www.geneontology.org/). The number of genes associated with each term was calculated, and hypergeometric testing was applied to identify GO terms significantly enriched in the differentially expressed gene set compared to the background proteome. GO terms with corrected P-value ≤ 0.05 were considered significantly enriched. Similarly, KEGG pathway enrichment analysis was performed using hypergeometric tests against the KEGG Pathway database. Pathways with Q-value ≤ 0.05 after multiple testing correction were defined as significantly enriched.

### Pseudotime and CytoTRACE analysis

For pseudotime trajectory analysis, we used both Monocle 2 and Monocle 3. In Monocle 2^81^, the gene expression matrix was used to order cells in pseudotime by projecting them onto a reduced dimensional space and reconstructing a tree-like trajectory. Cells with the smallest pseudotime value were defined as the most primitive population based on biological context. Genes whose expression significantly varied with pseudotime were identified using a smooth, nonlinear model, with FDR < 1×10⁻⁵ as the threshold for significance. In Monocle 3^82^, after preprocessing and PCA, nonlinear dimensionality reduction was performed using UMAP, followed by clustering. Partition-based graph abstraction (PAGA)^83^ was applied to identify differentiation partitions, grouping cell clusters with potential lineage relationships (FDR < 1%). Within each partition, trajectories were constructed and pseudotime values were calculated using the simplePPT graph embedding algorithm. Pseudotime values and partition information were then mapped back to the original Seurat UMAP embeddings for visualization. To assess differentiation potential, we performed CytoTRACE analysis^84^, which predicts cellular differentiation states based on gene count signatures. The CytoTRACE scores were mapped onto the Seurat UMAP embeddings to visualize the differentiation hierarchy of cell populations.

### RNA velocity analysis

RNA velocity analysis was performed to predict the developmental direction and transcriptional dynamics of cell populations. First, repeat regions in the *P. tomentosa* genome were annotated using RepeatMasker (v4.1.2) with the default repeat library to generate a repeat annotation file in GTF format. The BAM file generated by BSTMatrix from spatial transcriptome data was then sorted by cell barcode (CB tag) using samtools (v1.9)^85^ to produce a cell-sorted BAM file suitable for velocity analysis. Velocyto (v0.17.17)^86^ was applied to quantify spliced and unspliced transcripts. Using the cell-sorted BAM file, the gene annotation GTF file, and the repeat region GTF file as inputs, velocyto generated a loom count matrix containing spliced, unspliced, and ambiguous RNA abundances for each cell.

Downstream analysis was performed using scVelo (v0.2.4)^87,88^ in Python. The loom matrix was filtered and normalized with the following parameters: minimum shared counts of 20 per gene, selection of the top 2,000 highly variable genes, principal component analysis with 30 components, and construction of a neighborhood graph with 30 neighbors. The dynamical model implemented in scVelo was used to estimate the rates of transcription, splicing, and degradation, capturing the full kinetics of RNA metabolism. RNA velocity streams were visualized on the existing UMAP embeddings derived from snRNA-seq data, which had been annotated with cell type identities. Pseudotemporal ordering, driver gene identification, and PAGA trajectory inference were further performed to reconstruct the continuous differentiation paths and underlying regulatory dynamics.

### Weighted gene co-expression network analysis (WGCNA)

To identify co-expression modules associated with fiber cell development, we performed WGCNA^89^ using the WGCNA package in R. After filtering low-abundance and variably expressed gene, the normalized gene expression matrix was imported for network construction. A signed co-expression network was built using the automatic network construction function with default settings, and a soft-thresholding power of β = 5 was selected to achieve a scale-free topology fit. To identify modules of biological relevance, module–trait relationships were calculated by correlating module eigengenes with sample traits. For each module, intramodular connectivity was computed for all genes; genes with high connectivity were considered hub genes and potential key regulators.

For functional annotation, Gene Ontology (GO) and Kyoto Encyclopedia of Genes and Genomes (KEGG) pathway enrichment analyses were performed on genes within each module. Differentially expressed protein-coding genes were mapped to GO terms from the GO database (http://www.geneontology.org/). The number of genes associated with each term was counted. Hypergeometric tests were applied to identify terms significantly enriched compared to the background proteome, with P-values adjusted by Bonferroni correction (corrected P ≤ 0.05). For pathway analysis, hypergeometric tests against the KEGG Pathway database were performed, and pathways with Q ≤ 0.05 after multiple-testing correction were considered significantly enriched. The gene co-expression network was visualized using Cytoscape, based on node and edge files generated from hub genes identified in modules of interest.

### Phylogenetic analysis

Protein sequences of HD-Zip family members were retrieved from the Phytozome database (v14.0) and PlantTFDB 5.0. Multiple sequence alignments were performed using MEGA (v12.1), and a phylogenetic tree was constructed using the maximum likelihood method. The resulting tree was visualized and annotated using iTOL v7.

### Molecular cloning and binding assays

All genes were cloned from P. tomentosa cDNA libraries. Subcellular localization was performed by cloning coding sequences into pBI-121-EGFP vector, transforming into Agrobacterium GV3101, infiltrating *N. benthamiana* leaves, and observing EGFP fluorescence after 2 days. For protein-DNA interaction, Y1H assays were conducted by cloning promoter fragments into pABAi vector and integrating into Y1HGold genome, followed by transformation with pGADT7 constructs containing transcription factor coding sequences. For EMSA, recombinant proteins were expressed in *Escherichia coli* (*E. coli*), purified by affinity tags, and incubated with biotin-labeled L1-box probes; binding was detected by chemiluminescence. For dual-luciferase and GUS reporter assays, effector constructs (35S:*PtoPDF2*) and reporter constructs (target gene promoters driving LUC or GUS) were co-transformed into Agrobacterium and infiltrated into *N. benthamiana* leaves. LUC activity was measured after 2–3 days, and GUS activity was detected by histochemical staining or fluorometric assay. BiFC and LCA assays were performed using appropriate vectors, and fluorescence signals were observed under confocal microscopy. Data are presented as mean ± s.d. of three biological replicates. Statistical significance was determined by two-tailed Student’s t-test

## Data availability

The single-nucleus RNA sequencing and spatial transcriptomics data generated in this study have been deposited in the National Genomics Data Center (NGDC), China National Center for Bioinformation / Beijing Institute of Genomics, Chinese Academy of Sciences, under BioProject accession PRJCA060949 (https://ngdc.cncb.ac.cn/search/specific?db=bioproject&q=PRJCA060949). All other data supporting the findings of this study are available within the paper and its Supplementary Information, or from the corresponding author upon reasonable request.

## Acknowledgements

We thank our colleagues for helpful discussions during the course of this work. This work was supported by the National Natural Science Foundation of China (32672327, 31800555) and the National Key Research and Development Program of China (2024YFD2201004).

## Author contributions

Z.C. conceived the project, supervised the study and acquired funding. K.H. and H.W. contributed equally to this work. K.H. designed the study, performed the majority of the experiments and data analyses, and wrote the manuscript. H.W. participated in the experimental work and data analysis. J.T., X.Y. and T.Z. assisted with sample collection and data curation. X.A. and L.J. provided intellectual input on the experimental design and data interpretation. All authors discussed the results and commented on the manuscript, and Z.C. revised the manuscript. All authors read and approved the final manuscript.

## Supplementary information

**Supplementary Fig. 1.**
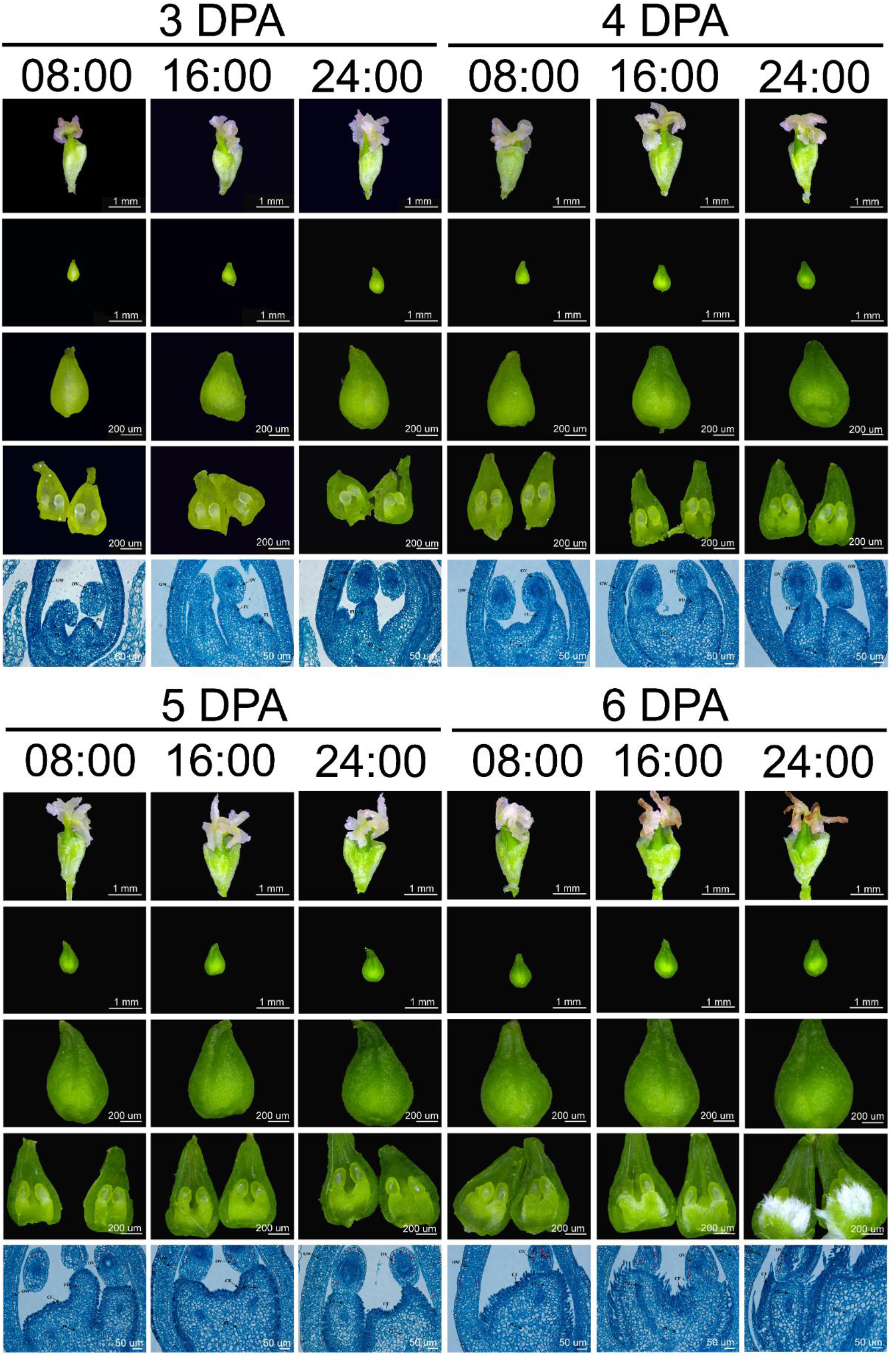
Morphological observations of poplar female capsule development from 3 to 6 DPA. Stereomicroscopic images showing the dynamic morphological changes of capsules at 3, 4, 5, and 6 days post-anthesis (DPA). The stigma transitions from pink to white and eventually to brown, while the capsule wall changes from yellow to dark green with increasing volume. Higher magnification views reveal the cellular differentiation processes within the ovary. Scale bars are indicated in the lower right corner of each image. OW, ovary wall; OV, ovule; FU, funiculus; PL, placenta; CF, catkin fibers.

**Supplementary Fig. 2.**
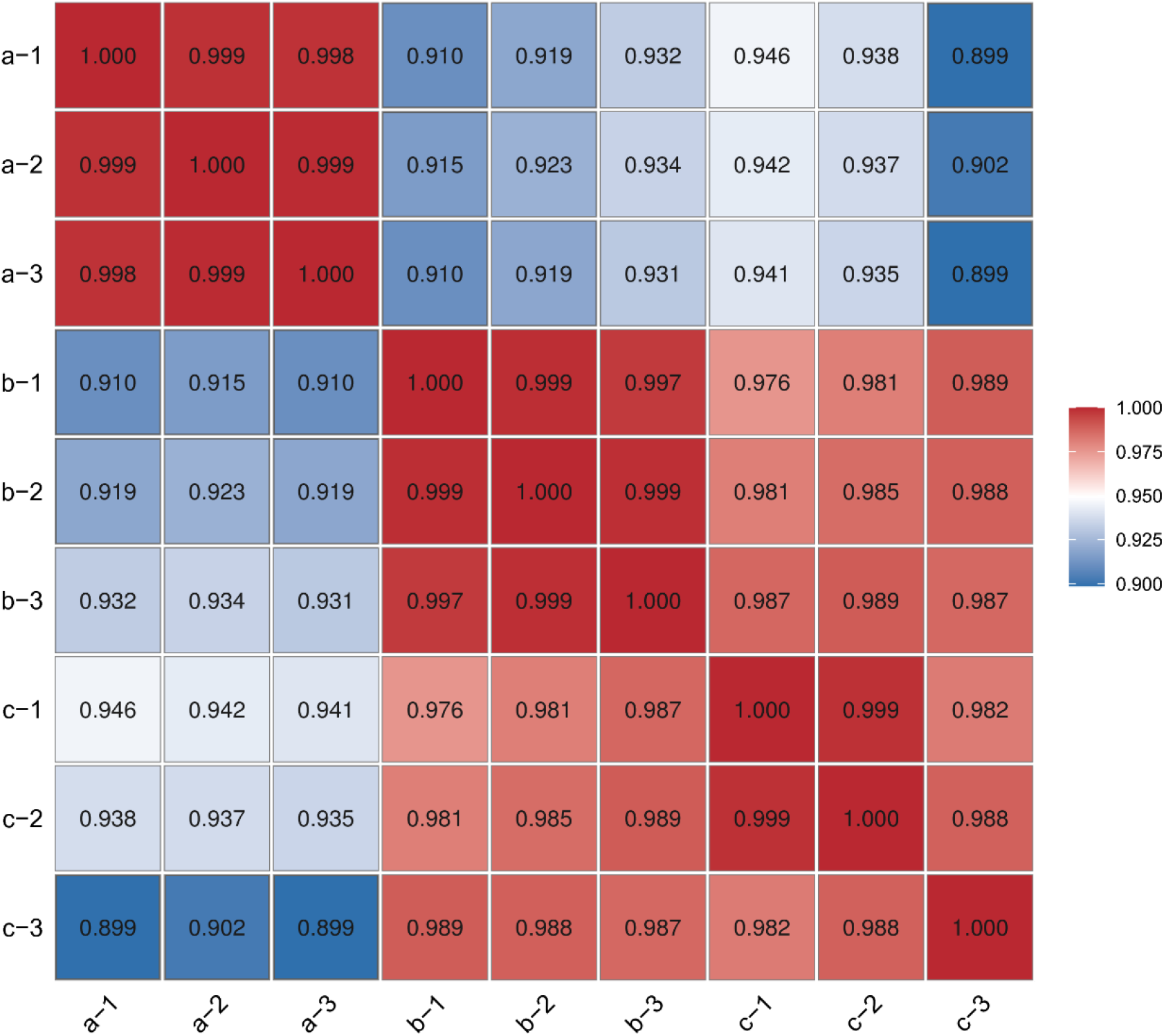
Heatmap of correlation analysis among samples. **a**, Stage 1 (4 DPA 24:00). b, Stage 2 (5 DPA 24:00). c, Stage 3 (6 DPA 16:00).

**Supplementary Fig. 3.**
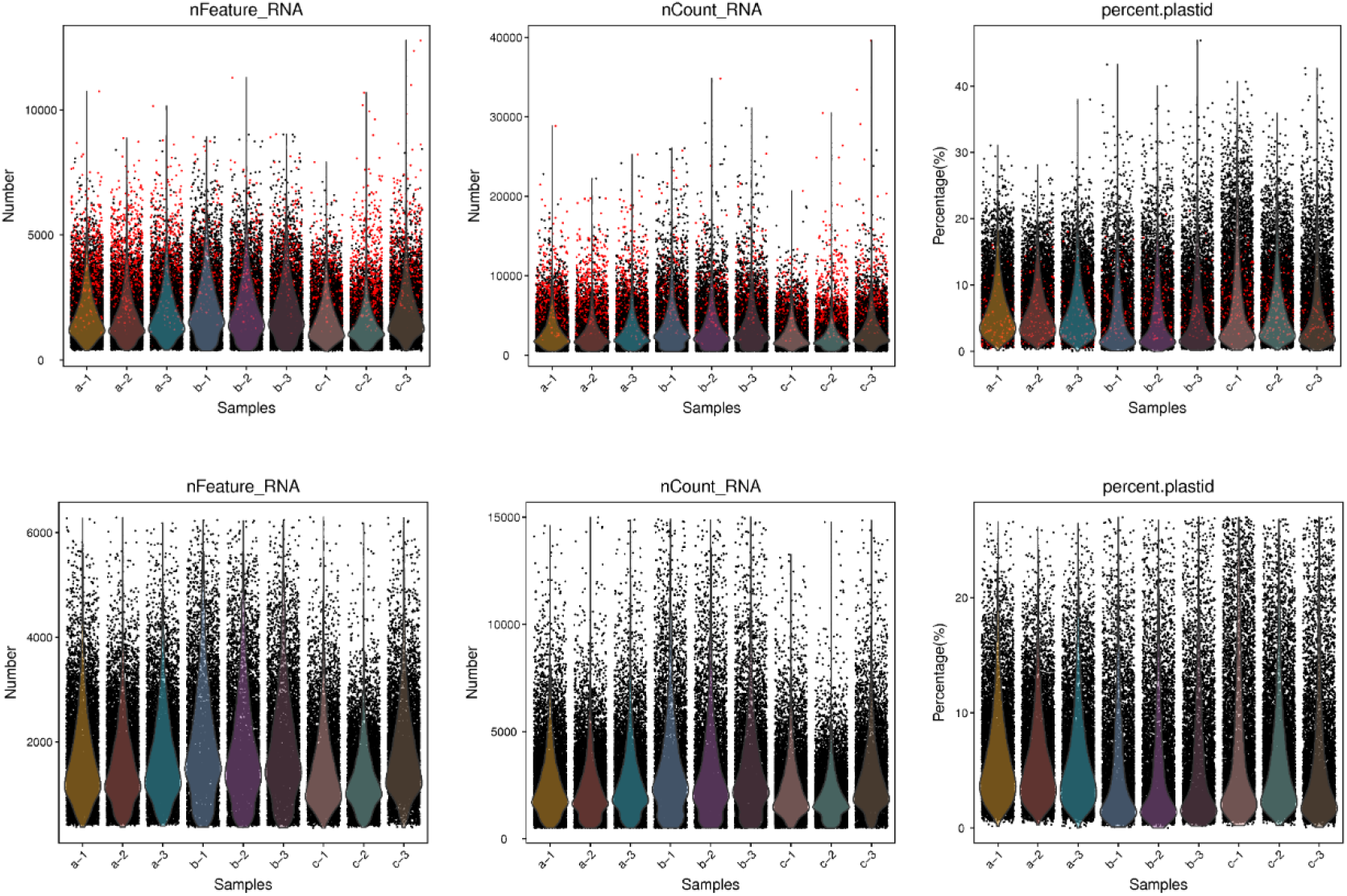
Violin plots of cellular data from nine samples before and after filtering. Upper panels: Distribution of basic cellular information across samples before filtering. Lower panels: Distribution of basic cellular information across samples after filtering. Left panels: Distribution of the number of genes detected per cell (Y-axis) for each sample. Middle panels: Distribution of the total UMI counts detected per cell (Y-axis) for each sample. Right panels: Distribution of the percentage of mitochondrial gene expression per cell (Y-axis) for each sample.

**Supplementary Fig. 4.**
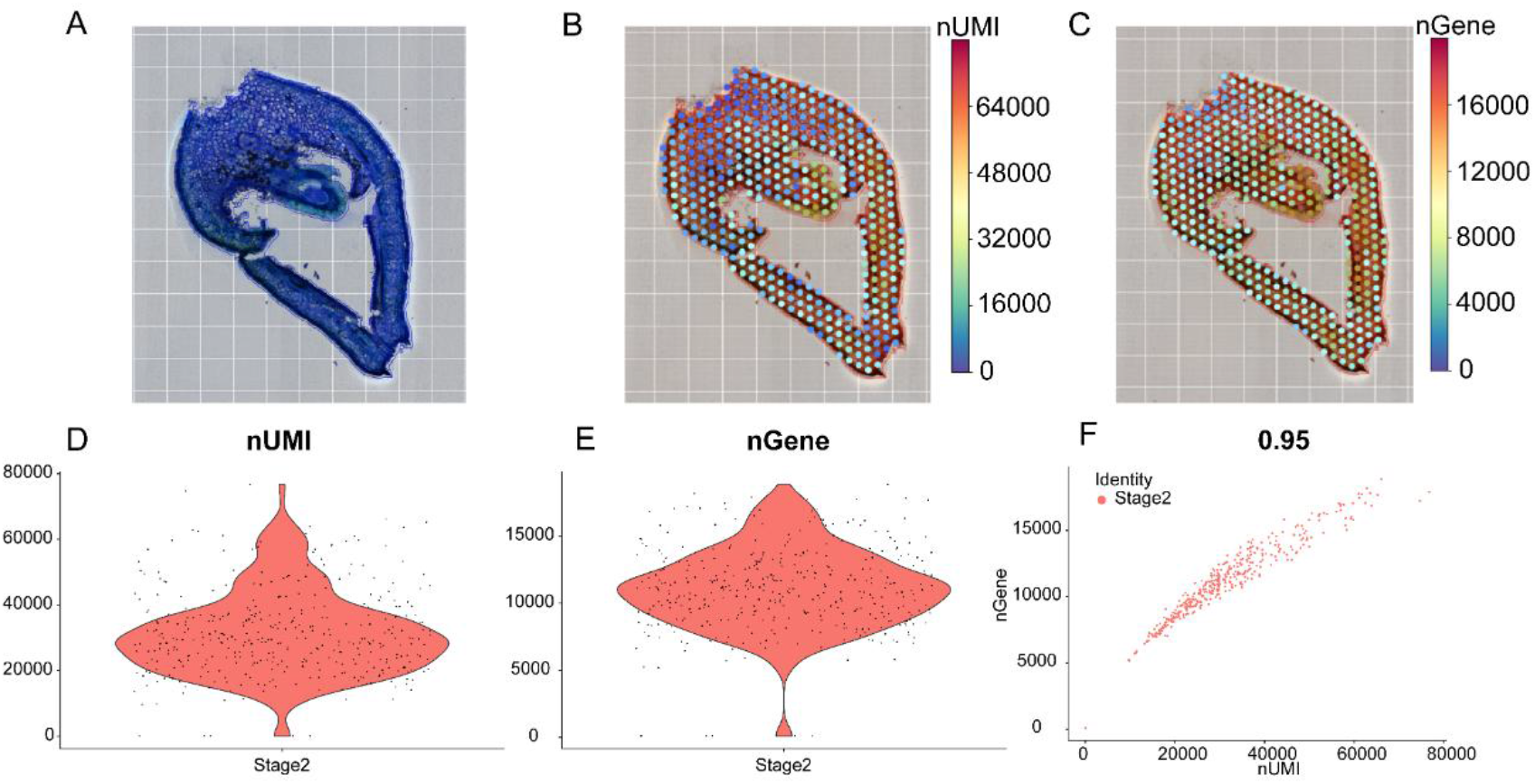
Quality control assessment of spatial transcriptome data. (**A**) Tissue section of a poplar female capsule at the second developmental stage (Stage 2) used for spatial transcriptome sequencing. (**B, C**) Basic quality control statistics showing the number of detected genes (nGene) and unique molecular identifiers (nUMI) per spot. The spatial distribution of UMI and gene counts across the tissue section is visualized, with each dot representing a single spatial spot. (**D, E**) Violin plots showing the distribution of UMI counts and gene counts per spot across samples. (**F**) Scatter plots showing the correlation between UMI counts and gene counts for each spot across samples.

**Supplementary Fig. 5.**
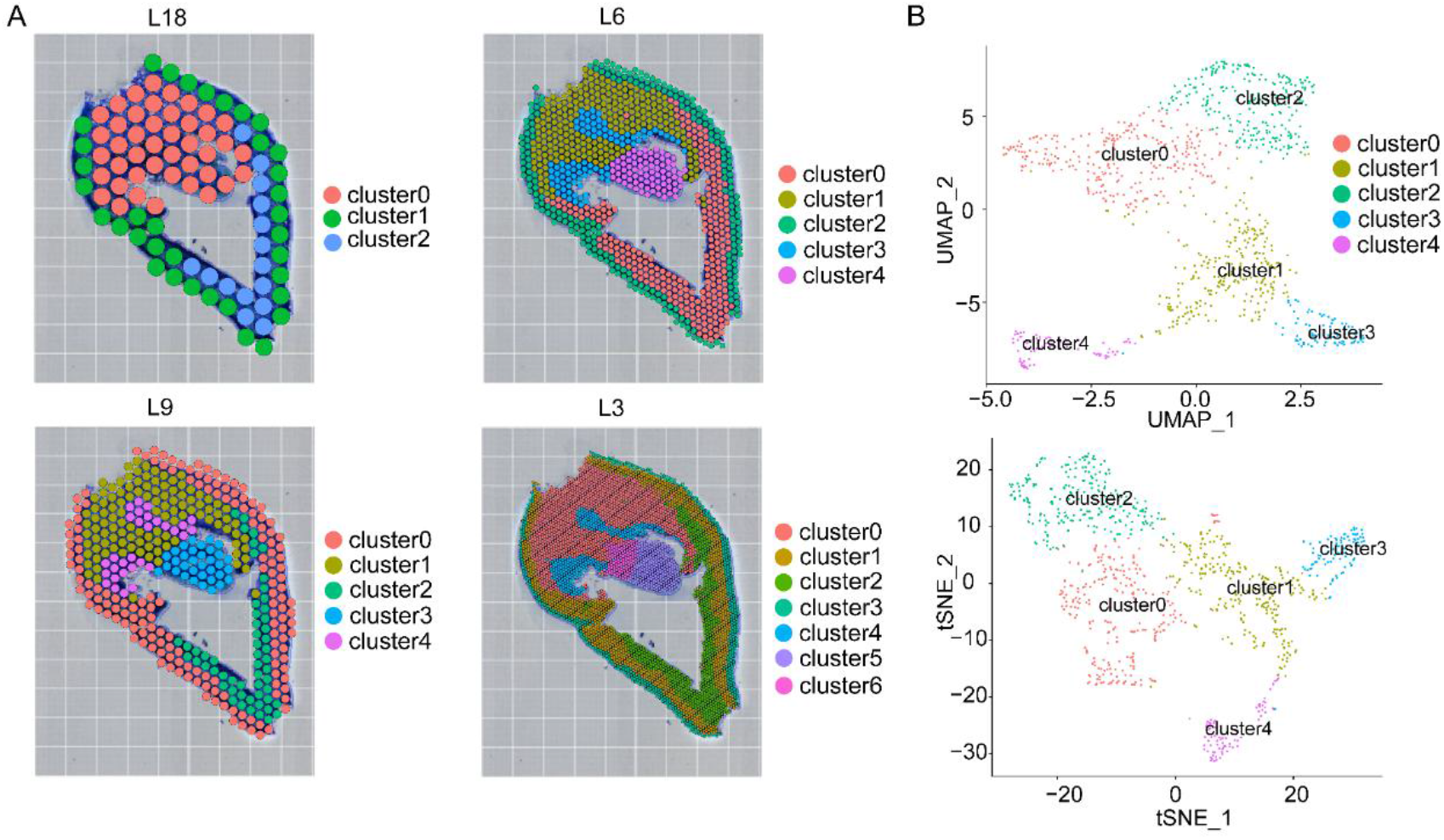
Visualization of spatial transcriptome clustering results at different levels. (**A**) Visualization of clustering results at different resolutions: level 18 (L18), L9, L6, and L3. (**B**) UMAP and t-Distributed Stochastic Neighborhood Embedding (t-SNE) visualization of cell clusters based on the L6 resolution. Cells were divided into five distinct clusters.

**Supplementary Fig. 6.**
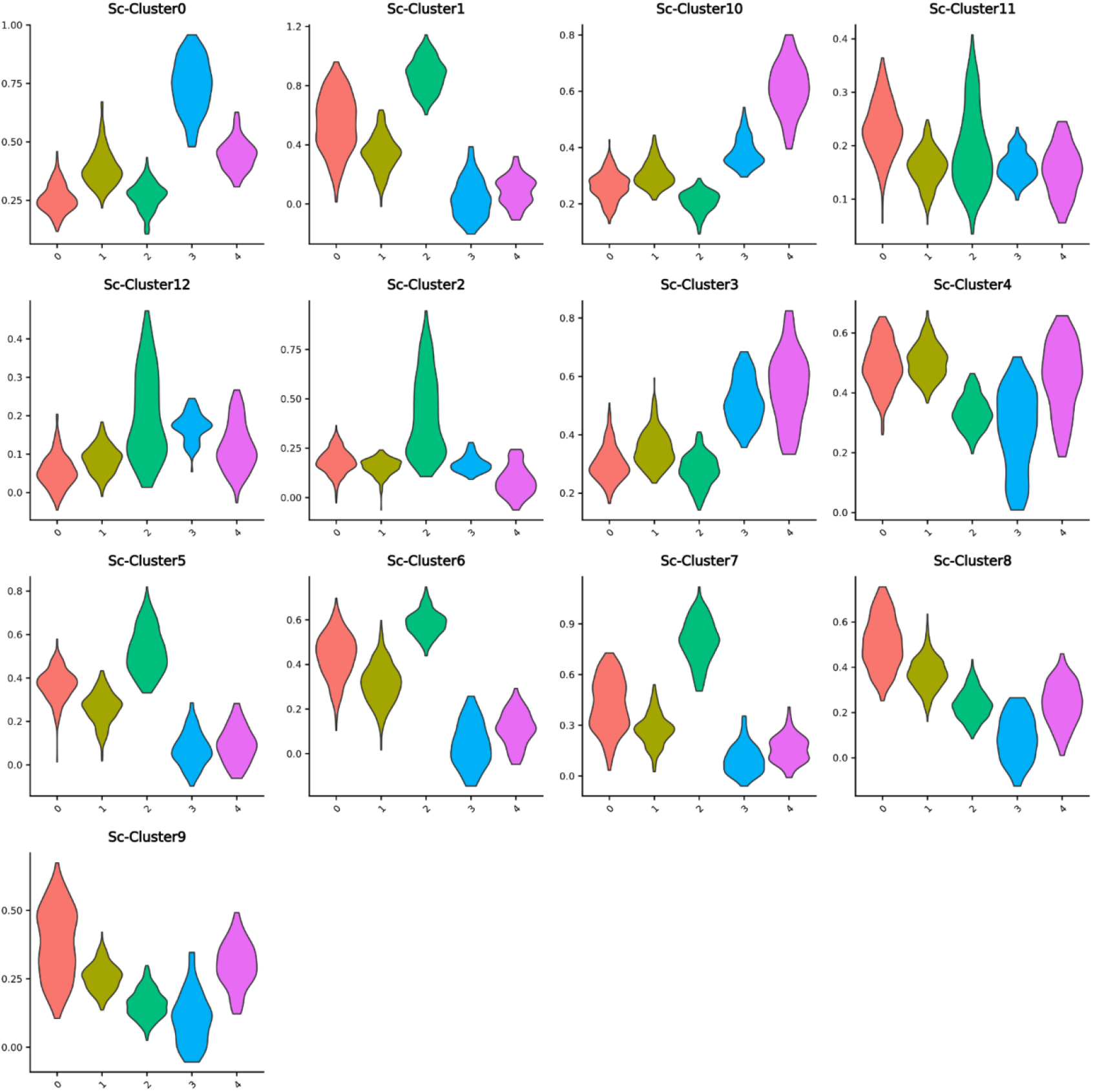
Violin plots showing the enrichment scores of marker gene sets from single-cell clusters in spatial transcriptome data. The x-axis represents distinct spatial tissue regions, and the y-axis represents the calculated enrichment score. For each target gene set, the enrichment score is defined as the average expression of the target genes minus the average expression of a background gene set.

**Supplementary Fig. 7.**
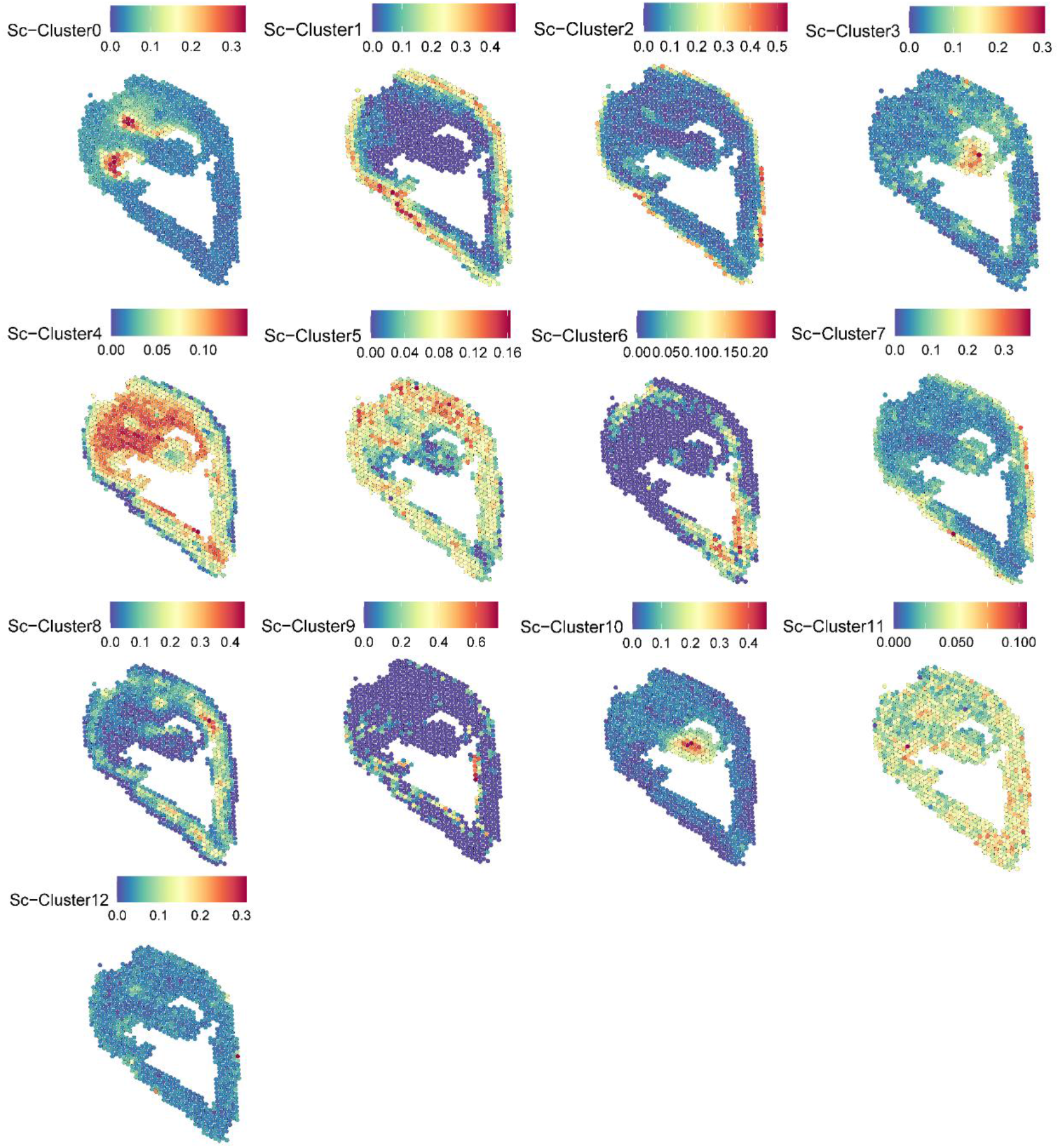
Spatial mapping of single-cell transcriptome-derived cell clusters. Spatial expression patterns of individual cell clusters (clusters 0 to 12) are shown, with colors ranging from blue to red, where red indicates higher expression levels.

**Supplementary Fig. 8.**
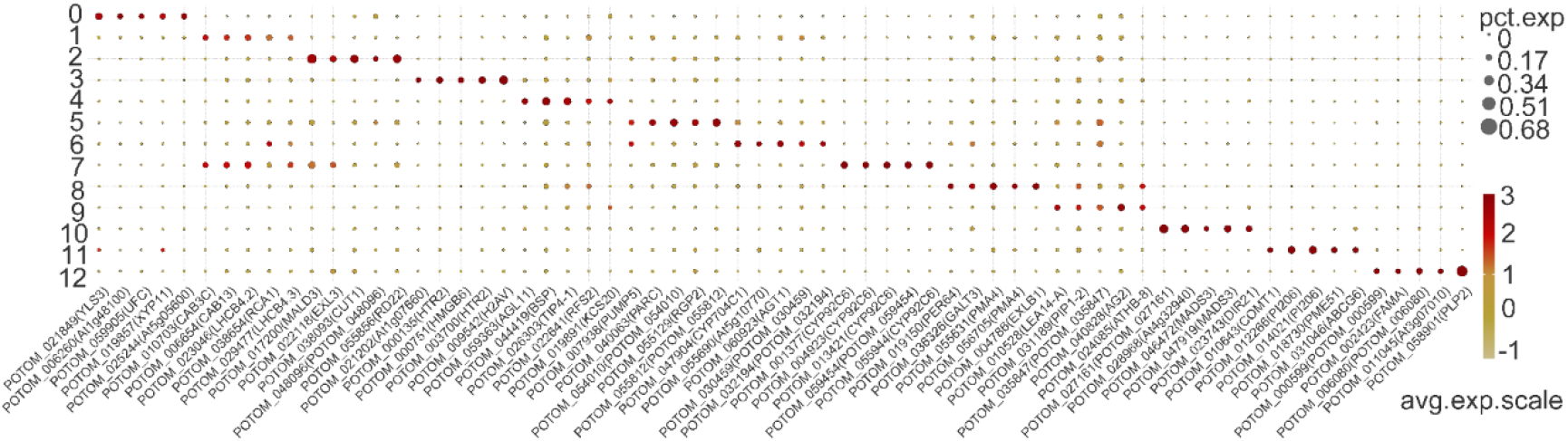
Bubble plot showing specifically highly expressed marker genes across different single-cell clusters.

**Supplementary Fig. 9.**
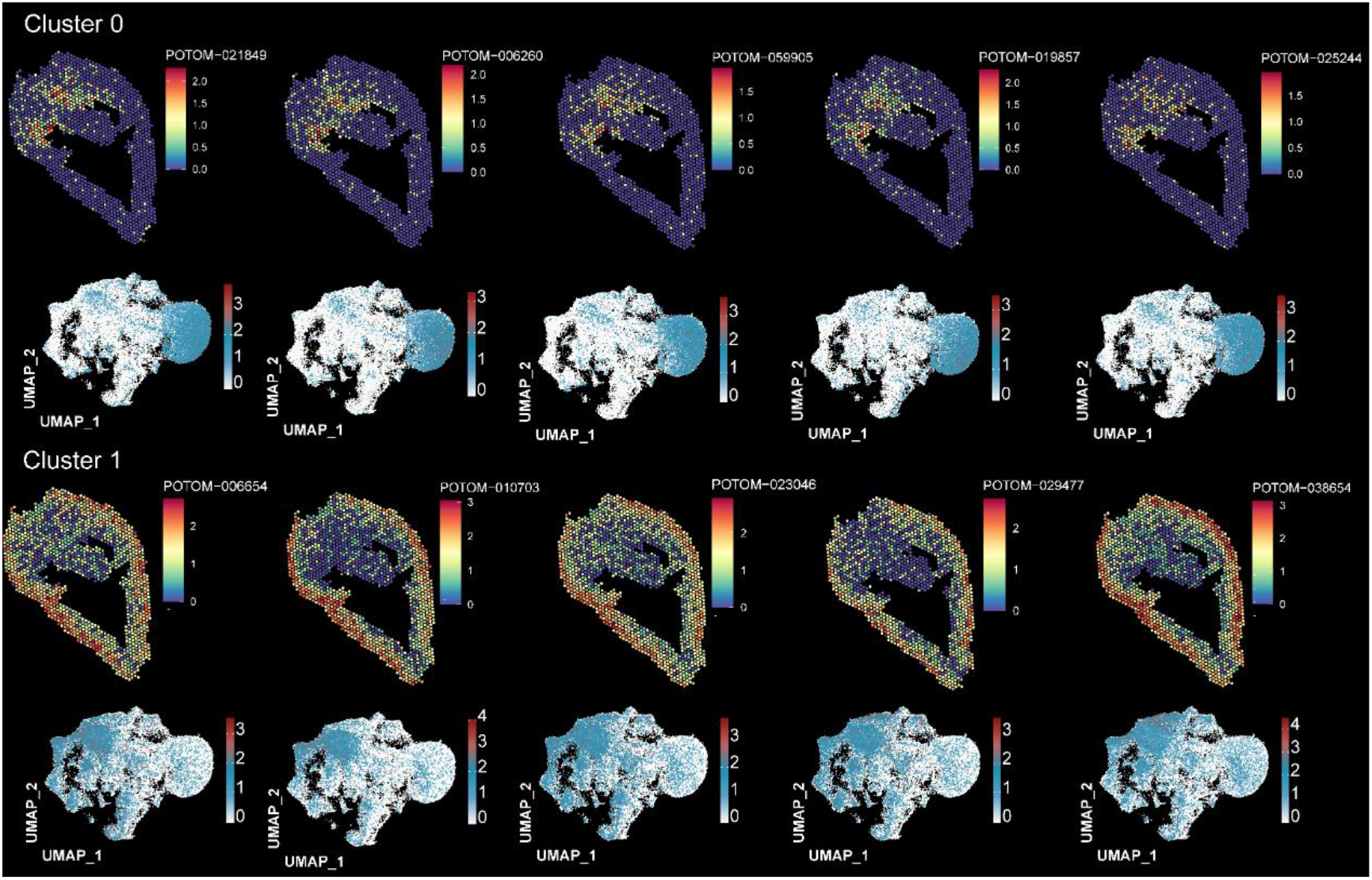
Spatial expression patterns of marker genes for clusters 0 and 1.

**Supplementary Fig. 10.**
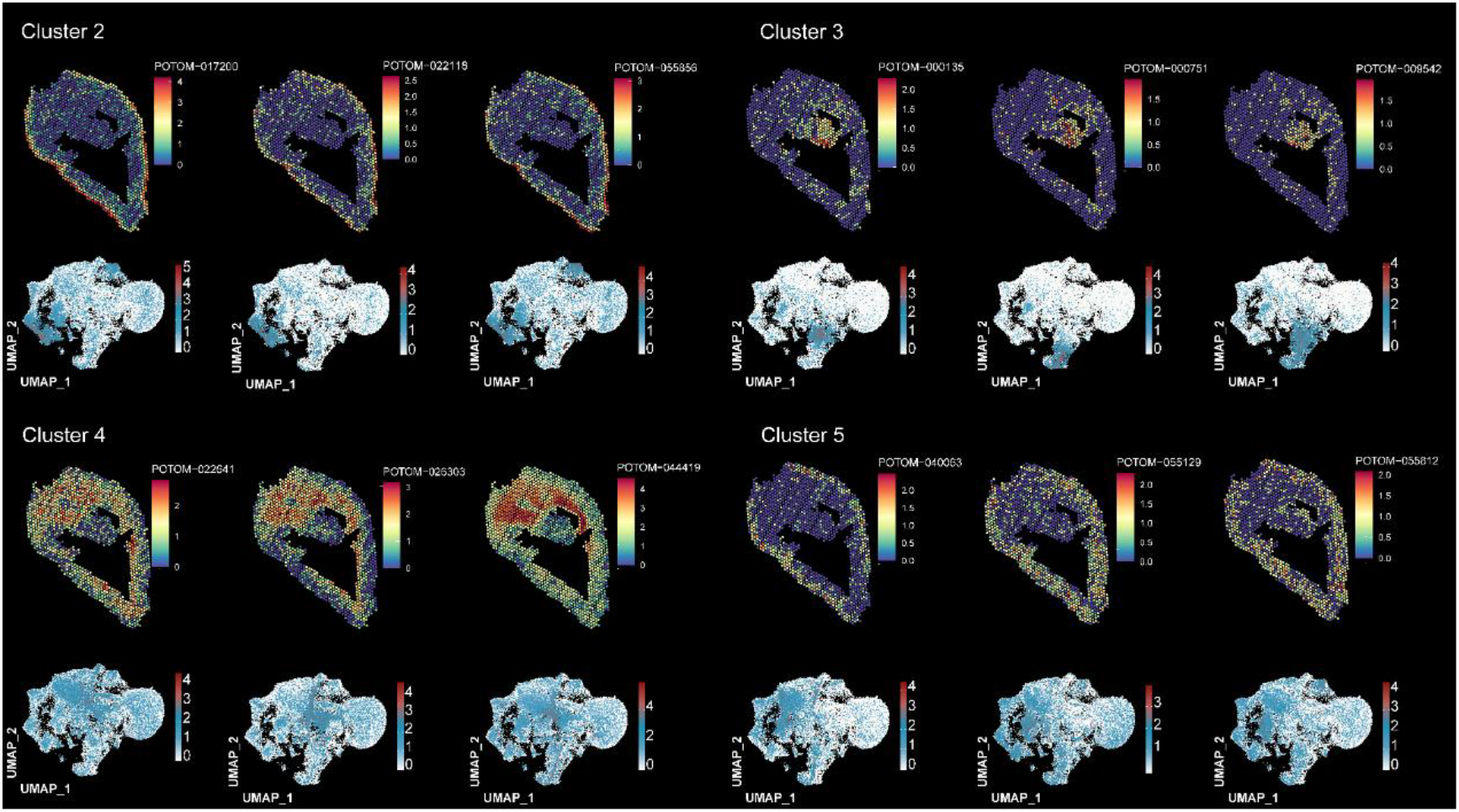
Spatial expression patterns of marker genes for clusters 2, 3, 4, and 5.

**Supplementary Fig. 11.**
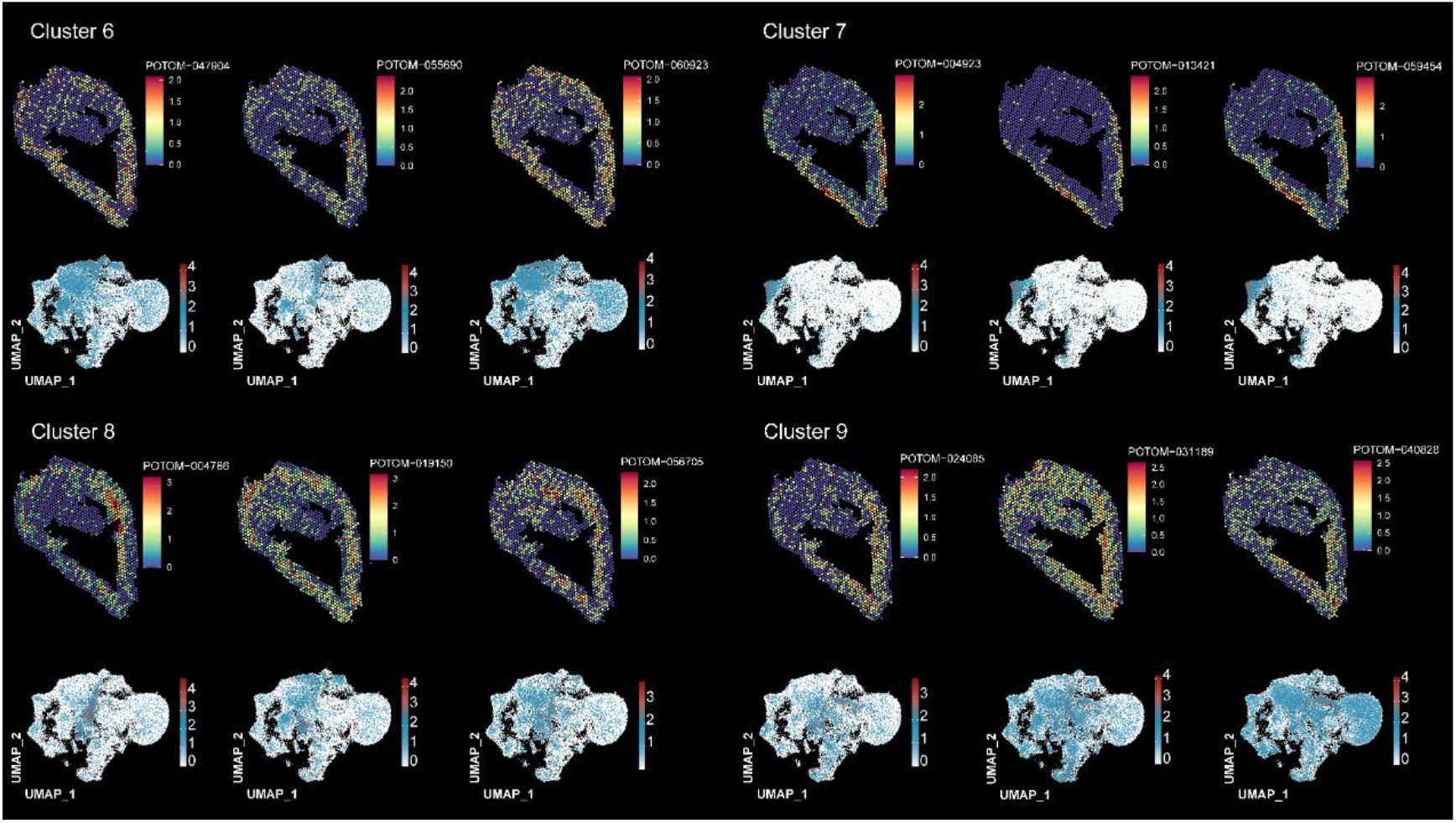
Spatial expression patterns of marker genes for clusters 6, 7, 8, and 9.

**Supplementary Fig. 12.**
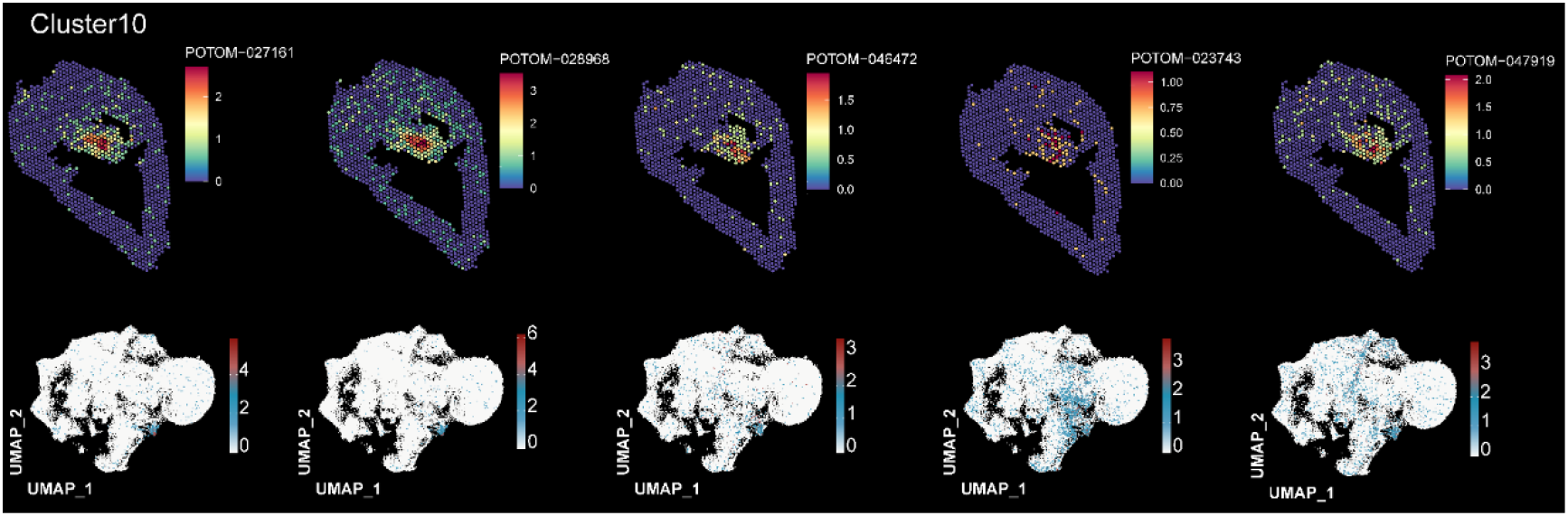
Spatial expression patterns of marker genes for clusters 10.

**Supplementary Fig. 13.**
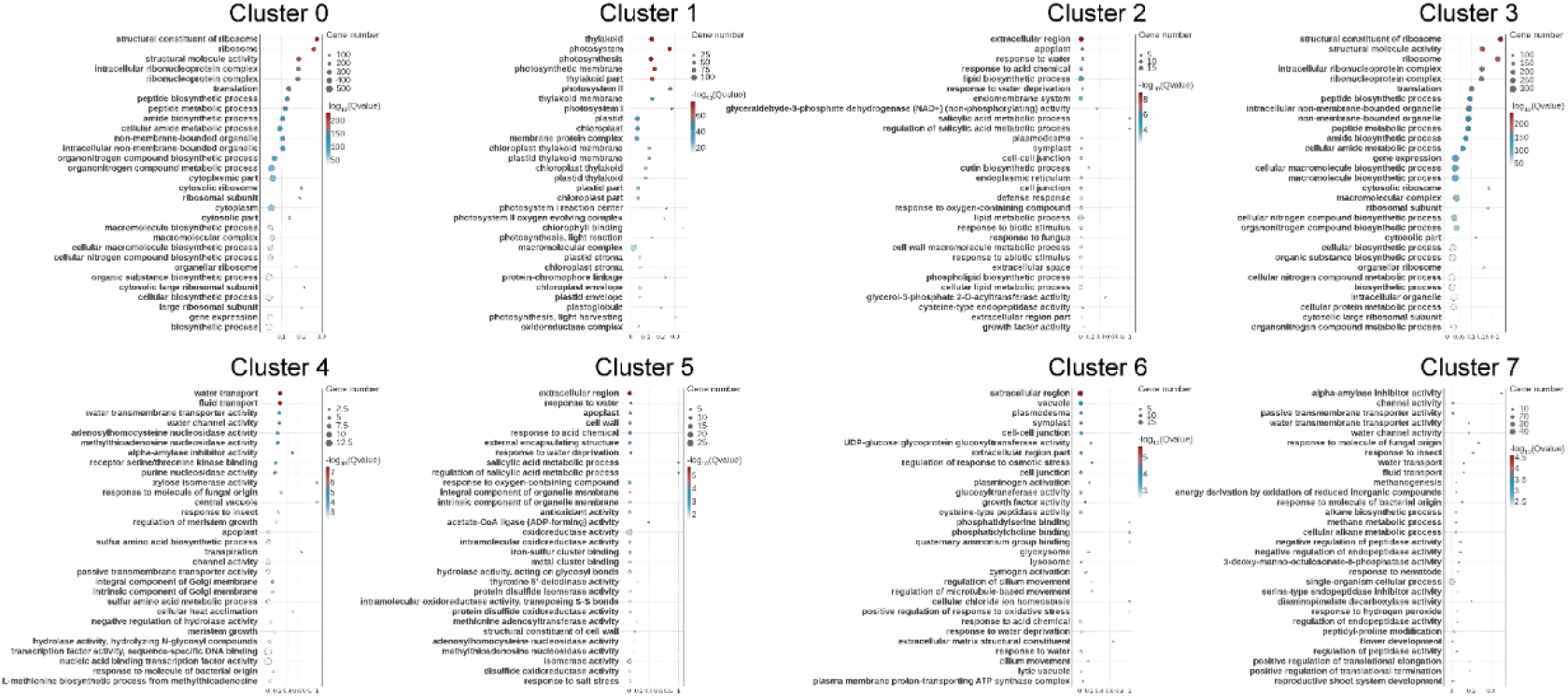
GO enrichment results for clusters 0 through 7.

**Supplementary Fig. 14.**
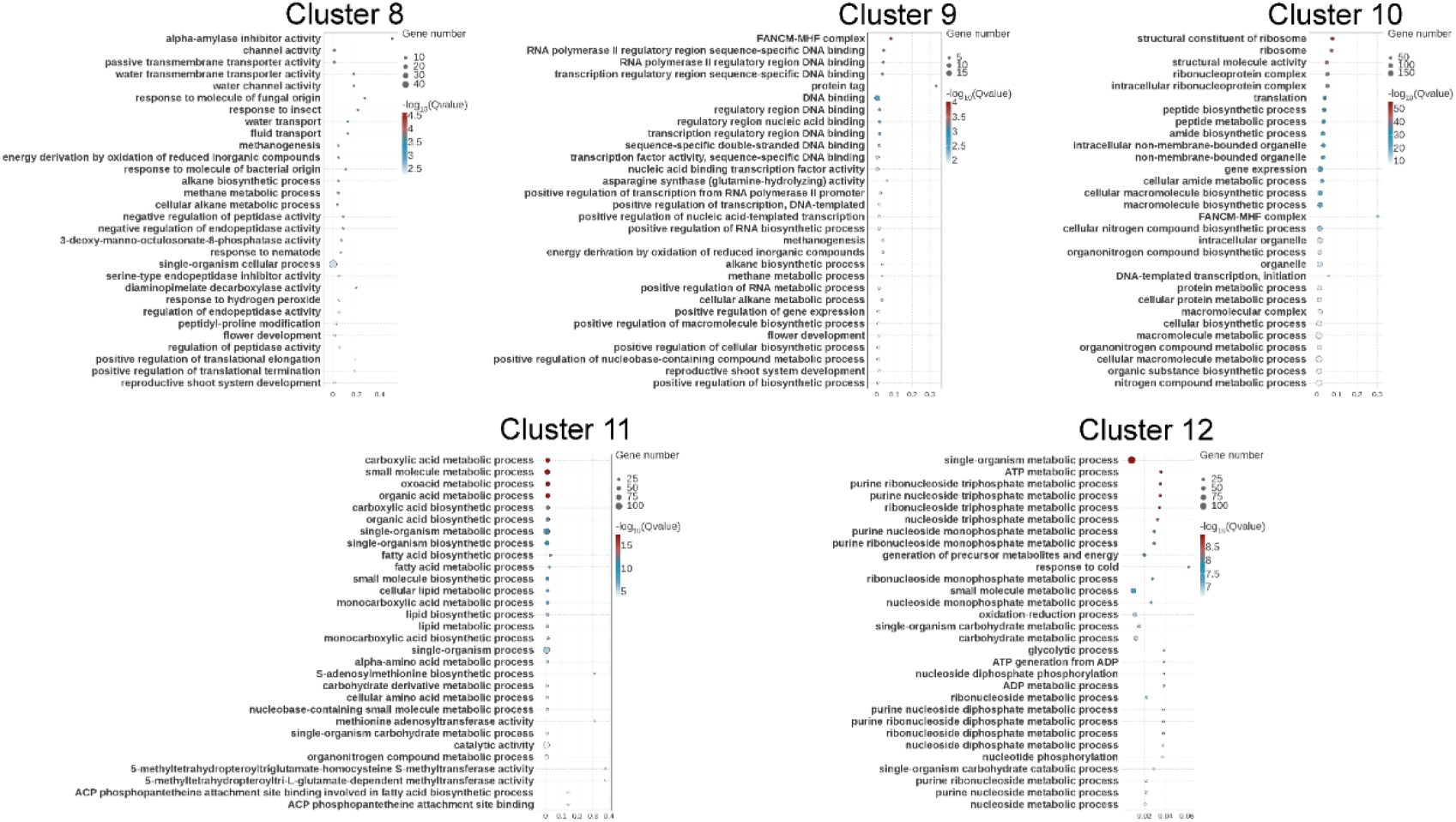
GO enrichment results for clusters 8 through 12.

**Supplementary Fig. 15.**
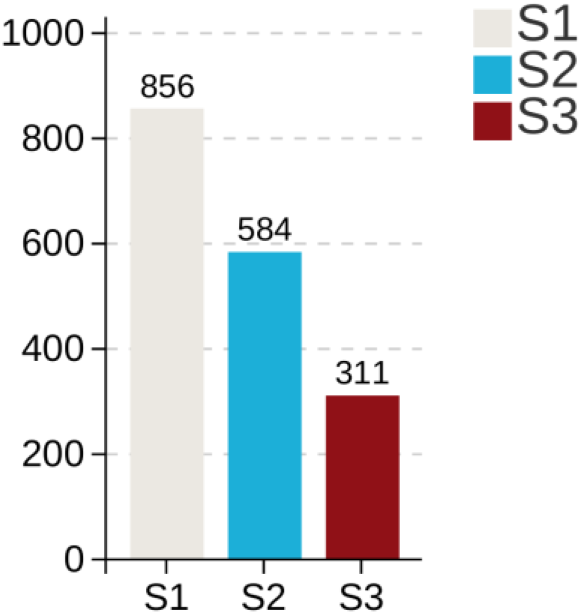
Bar plots showing the number of upregulated genes in fiber cells (cluster 0) at each of the three developmental stages. S1, stage 1; S2, stage 2; S3, stage 3.

**Supplementary Fig. 16.**
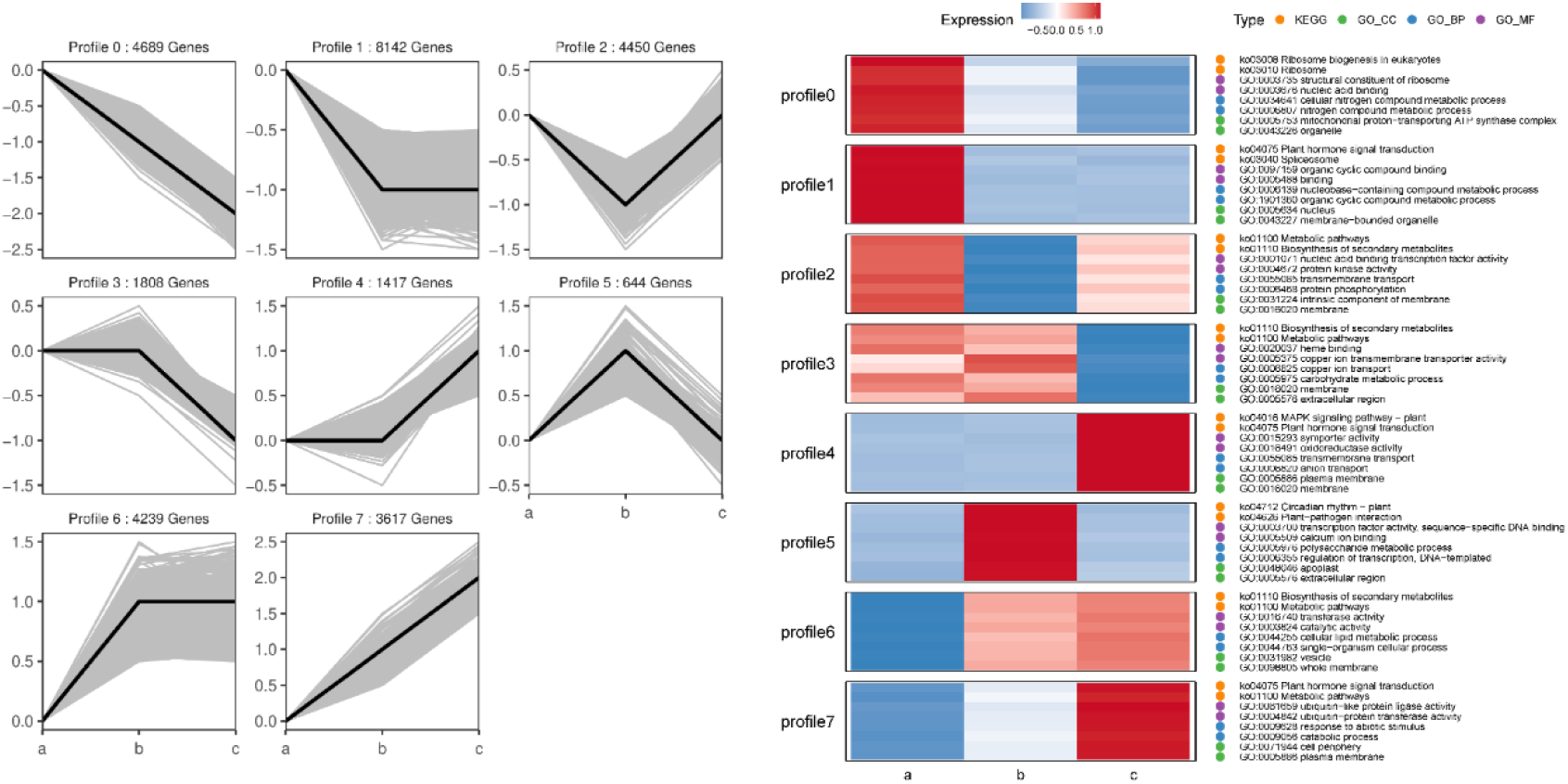
Trend analysis of all genes expressed in fiber cells across the three developmental stages. The left panel shows the eight distinct expression patterns identified, with the number of genes assigned to each trend indicated. The right panel displays the significantly enriched functional pathways (KEGG and GO terms) associated with each expression trend.

**Supplementary Fig. 17.**
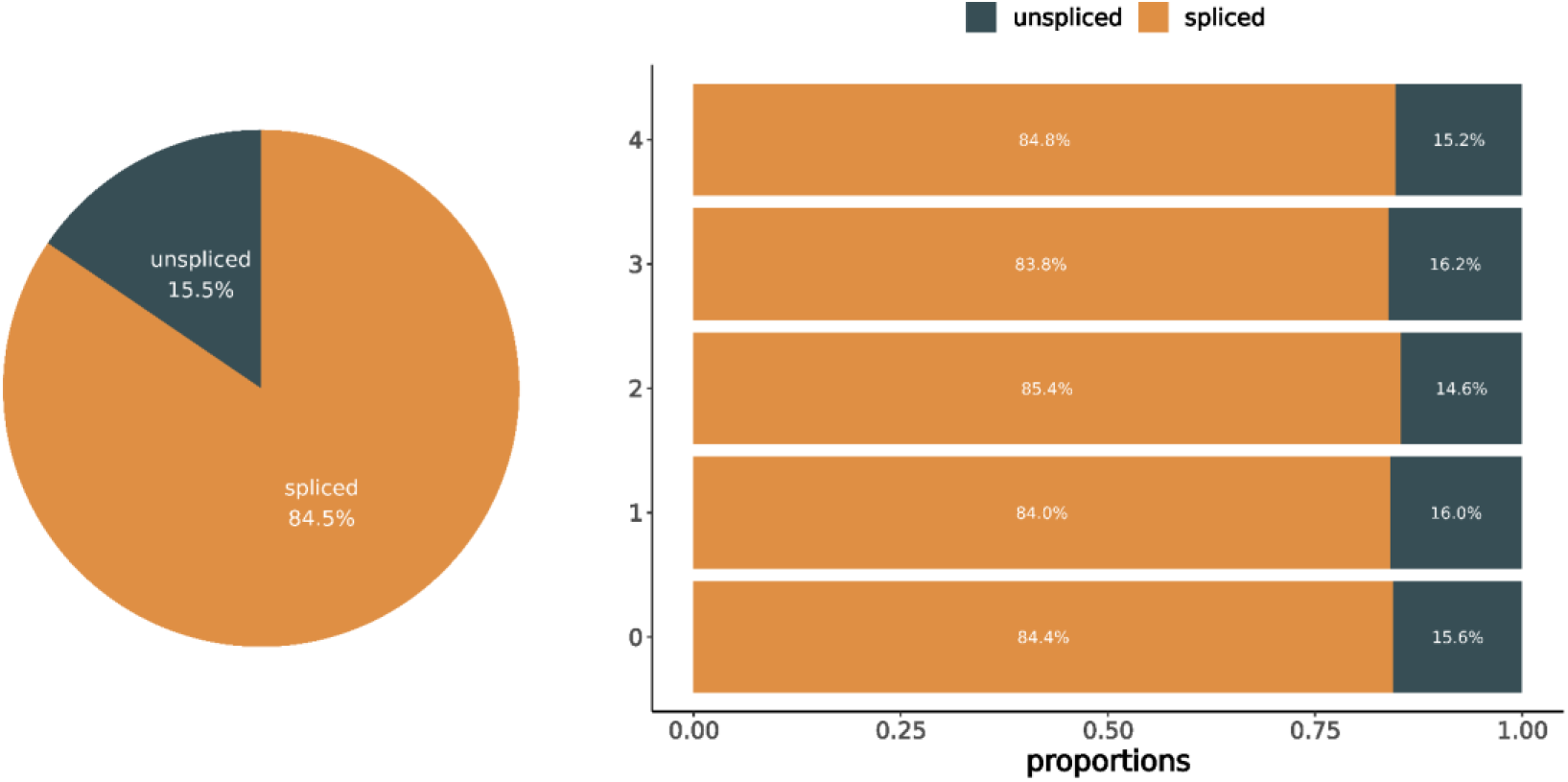
Proportions of unspliced (nascent) and spliced (mature) mRNA abundance. For sample, the abundance of unspliced and spliced mRNA was quantified in high-quality cells, and the ratio of unspliced to spliced mRNA abundance was calculated for each cell type. The y-axis represents cell types, and the x-axis represents the abundance ratio of unspliced (nascent) mRNA (green) to spliced (mature) mRNA (yellow).

**Supplementary Fig. 18.**
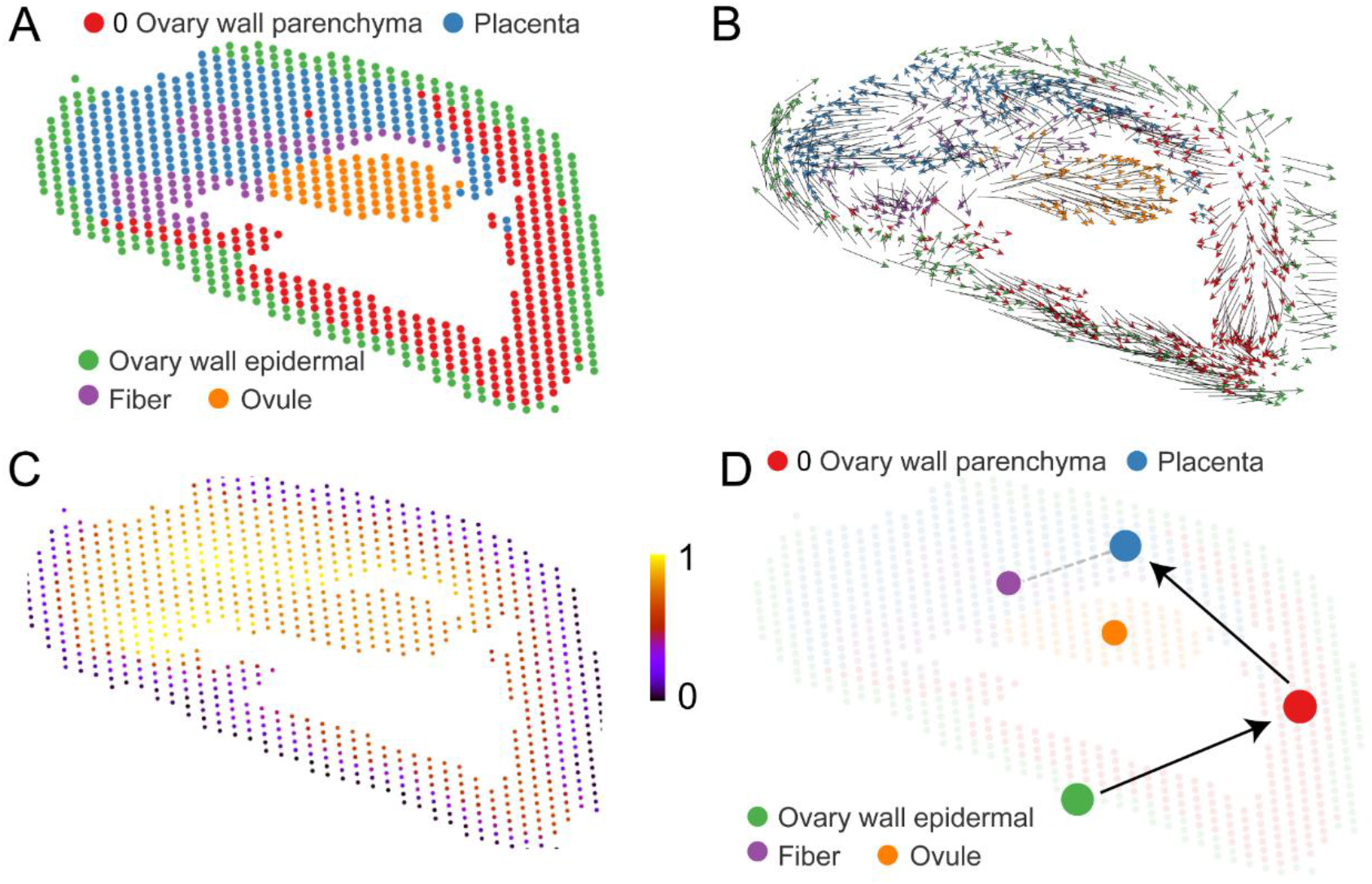
RNA velocity analysis. (**A**) Spatial transcriptome mapping of poplar female capsules, revealing five distinct cell types, each represented by a different color. (**B**) RNA velocity stream plot. Each dot represents a single cell, with cells of the same cell type shown in the same color. Arrows indicate the direction and magnitude of RNA velocity for individual cells. (**C**) Latent time inferred from RNA velocity. Each dot represents a single cell, with darker colors indicating cells closer to the onset of differentiation and lighter colors representing cells further along the differentiation trajectory. (**D**) PAGA (Partition-based Graph Abstraction) analysis based on RNA velocity. Each dot represents a single cell, with cells of the same cell type shown in the same color. Arrows indicate potential differentiation directions among cells.

**Supplementary Fig. 19.**
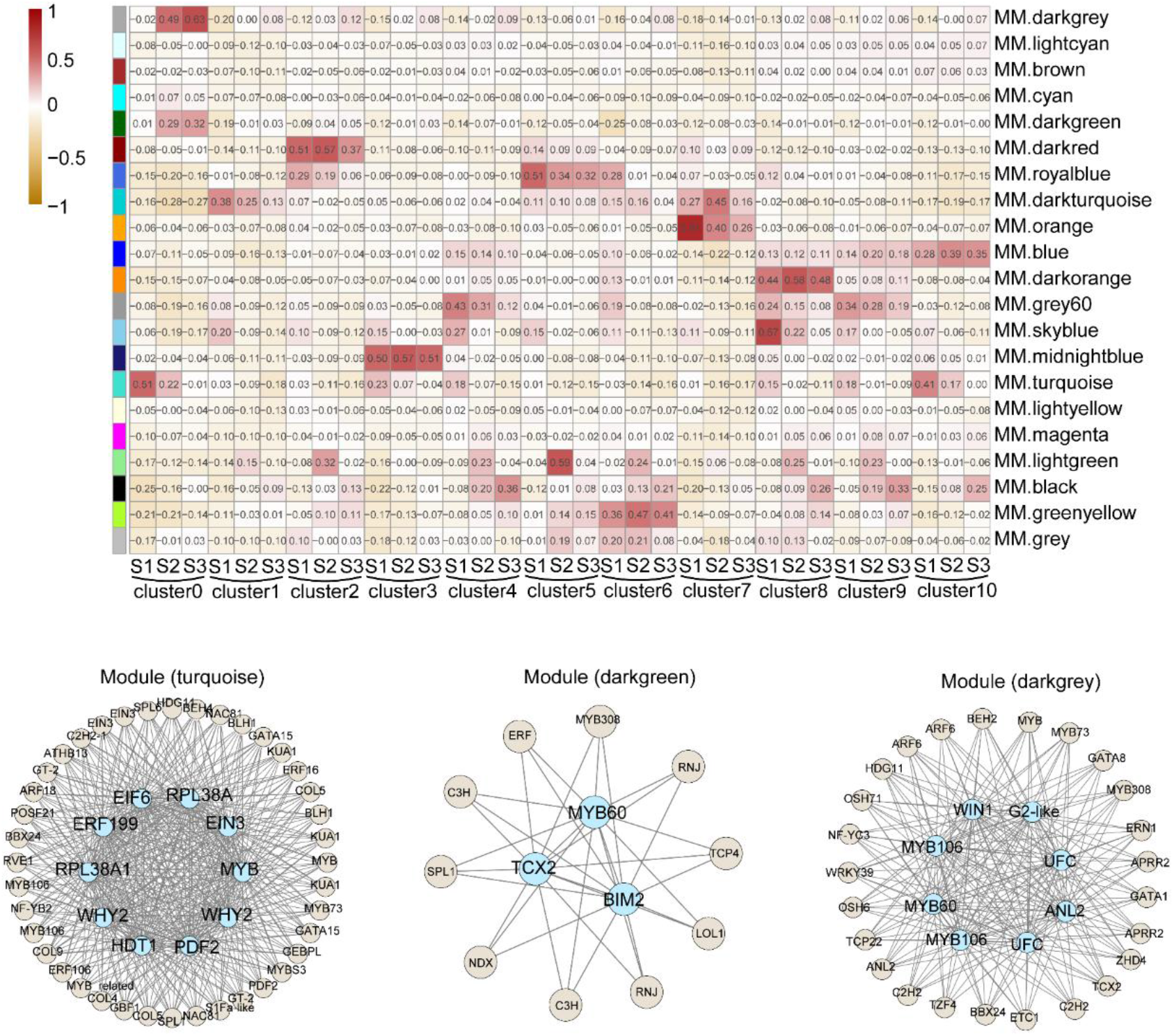
WGCNA and gene network construction based on snRNA-seq data. Upper panels: WGCNA identified 21 distinct modules, with increasing red intensity indicating stronger correlation. Lower panels: Gene regulatory networks constructed for the turquoise, darkgreen, and darkgrey modules. Hub transcription factors with high edge weights are positioned at the center of each network.

**Supplementary Fig. 20.**
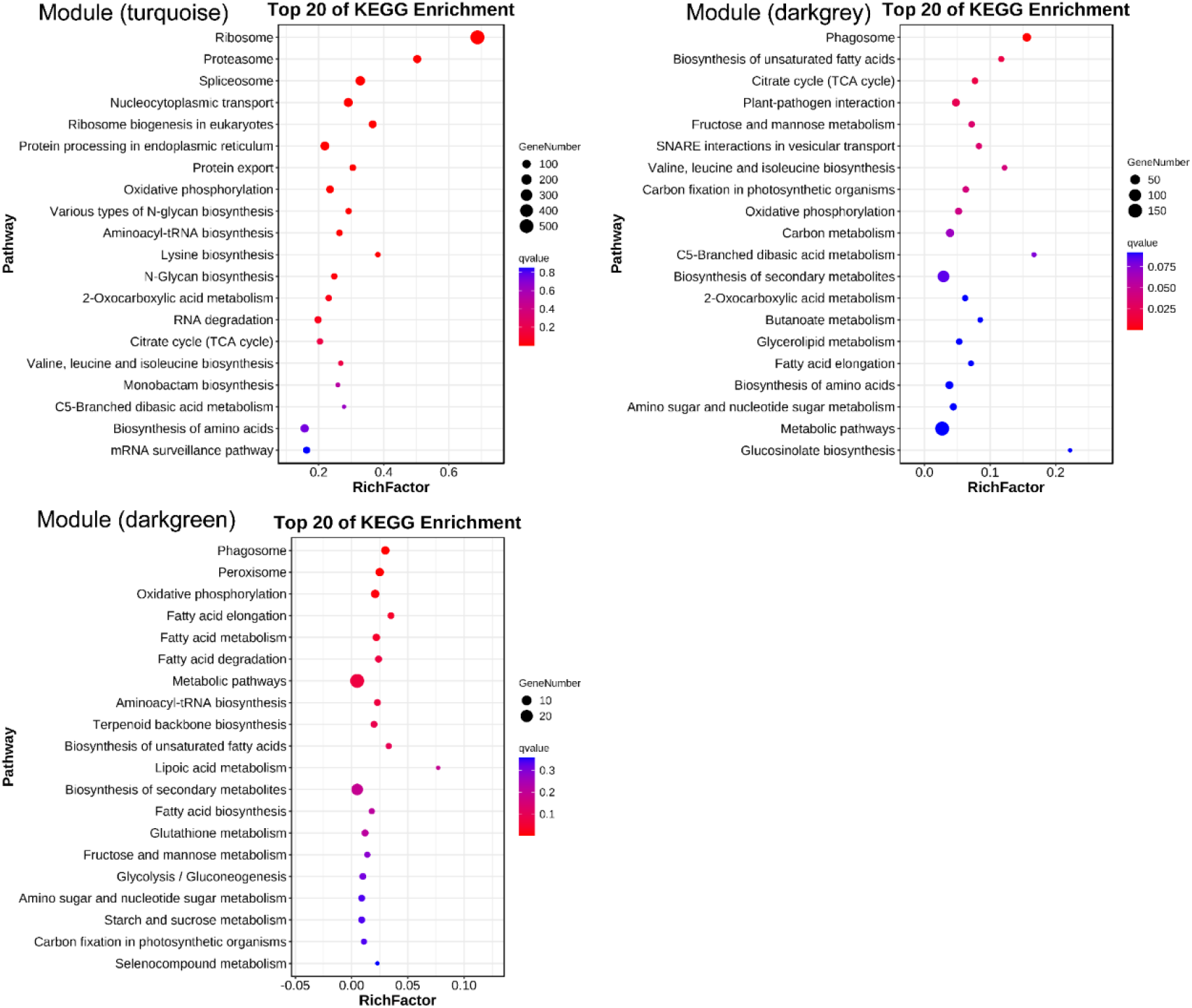
Top 20 enriched KEGG pathways in the turquoise, darkgreen, and darkgrey modules.

**Supplementary Fig. 21.**
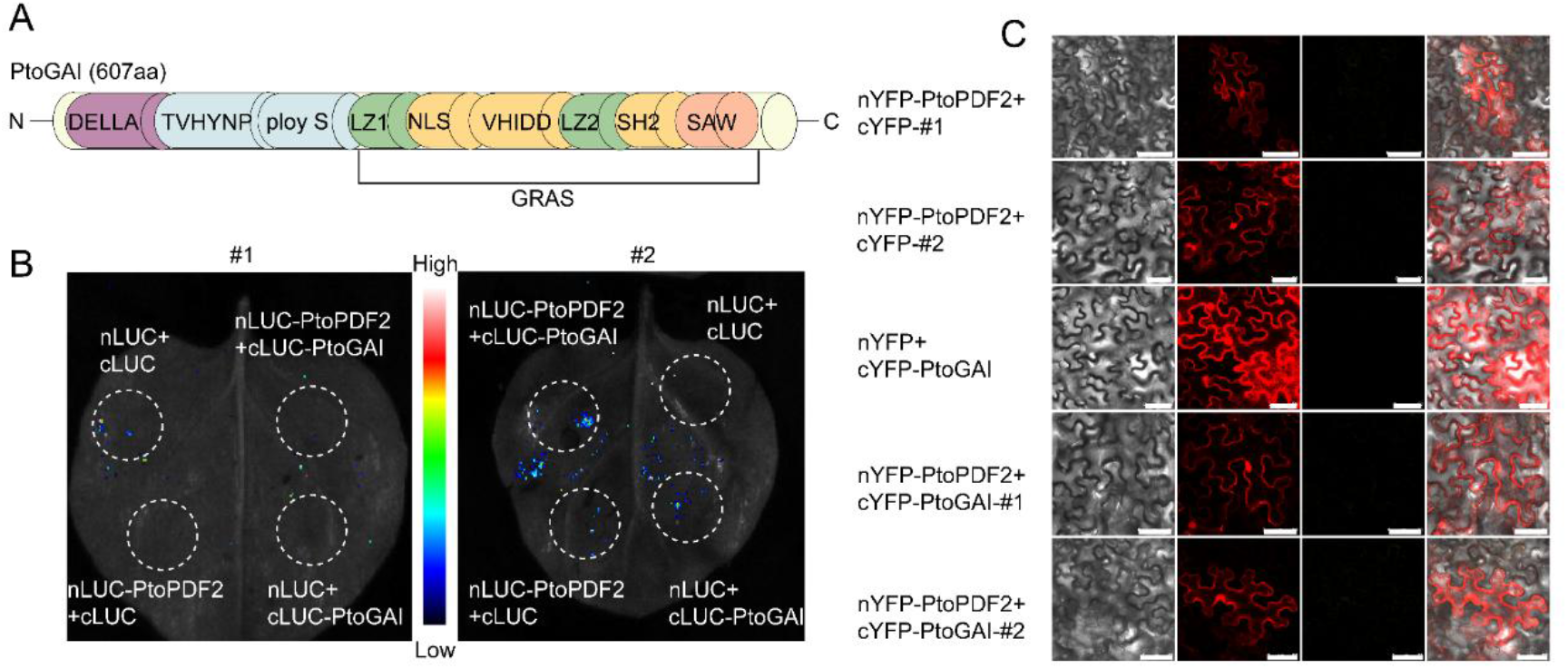
Interaction analysis of PtoGAI. (**A**) Schematic diagram of PtoGAI structure depicting the conserved domains characteristic of DELLA family proteins. (**B**) Luciferase complementation assays (LCA) in *N. benthamiana* showing no detectable in vivo interaction between either PtoPDF2 variant and PtoGAI. (**C**) Bimolecular fluorescence complementation (BiFC) assays confirming the absence of in vivo interaction. Scale bars, 75 μm (top row) and 50 μm (others).

## Description of Additional Supplementary Files

### Supplementary Data Legends

Supplementary Data 1. Gene Ontology (GO) enrichment pathways of the 13 cell clusters identified in this study.

Supplementary Data 2. Commonly upregulated differentially expressed genes in fiber cells across three developmental stages.

Supplementary Data 3. Commonly downregulated differentially expressed genes in fiber cells across three developmental stages.

Supplementary Data 4. Commonly regulated differentially expressed genes in fiber cells across three developmental stages.

Supplementary Data 5. The 49 core transcription factors identified among the commonly regulated genes across three developmental stages of fiber cells.

Supplementary Data 6. Gene Ontology (GO) enrichment analysis of stage-specific upregulated genes in fiber cells across three developmental stages.

Supplementary Data 7. Gene Ontology (GO) enrichment pathways of three gene modules identified along the pseudotemporal trajectory of fiber cell development.

Supplementary Data 8. Gene Ontology (GO) enrichment pathways of the three fiber cell subtypes.

Supplementary Data 9. Gene lists of the three fiber cell subtypes.

Supplementary Data 10. KEGG pathway enrichment analysis of three WGCNA modules associated with fiber cell development.

Supplementary Data 11. List of genes contained in the three WGCNA modules associated with fiber cell development.

