## Supplementary Table for "Single-cell spatiotemporal transcriptomics reveals the origin, cellular heterogeneity and regulatory network of seed fiber cells in poplar"

Supplementary Table. 1

Quality control of single-cell sequencing data

| Samples | before_filter_num | after_filter_num | pct | before_filter_median_UMI_per_cell | after_filter_median_UMI_per_cell | before_filter_median_genes_per_cell | after_filter_median_genes_per_cell | before_filter_median_Plastid_per_cell | after_filter_median_Plastid_per_cell |
| --- | --- | --- | --- | --- | --- | --- | --- | --- | --- |
| a-1 | 12449 | 11238 | 90.27% | 2362 | 2198 | 1579 | 1489 | 5.065234 | 5.25960668 |
| a-2 | 14151 | 12611 | 89.12% | 2170 | 2021 | 1490 | 1392 | 4.809725 | 4.898336414 |
| a-3 | 11327 | 10324 | 91.15% | 2453 | 2327.5 | 1628 | 1555 | 4.694836 | 4.791588684 |
| b-1 | 8551 | 7884 | 92.20% | 2953 | 2785 | 1853 | 1761.5 | 3.435469 | 3.282439978 |
| b-2 | 8939 | 8236 | 92.14% | 2715 | 2593 | 1731 | 1658 | 3.389831 | 3.194978218 |
| b-3 | 8392 | 7741 | 92.24% | 2844.5 | 2707 | 1779 | 1705 | 3.696758 | 3.582838886 |
| c-1 | 10186 | 9218 | 90.50% | 2187.5 | 2080.5 | 1376 | 1319 | 4.635477 | 4.359423843 |
| c-2 | 12432 | 11200 | 90.09% | 1877 | 1773 | 1243 | 1180 | 4.584041 | 4.573991179 |
| c-3 | 9353 | 8588 | 91.82% | 2651 | 2532.5 | 1698 | 1625 | 3.636364 | 3.672160513 |

Supplementary Table. 2

1. Quality assessment of raw spatial transcriptome data

| sampleID | ReadSum | BaseSum | GC(%) | N(%) | Q20(%) | Q30(%) |
| --- | --- | --- | --- | --- | --- | --- |
| Stage2 | 1168941080 | 3.5068E+11 | 3392.00% | 0.02 | 95.76 | 91.56 |

2. Alignment of spatial transcriptome data to the reference genome

| sampleID | Reads Mapped to Genome | Reads Mapped Confidently to Genome | Reads Mapped Confidently to Intergenic Regions | Reads Mapped Confidently to Intronic Regions | Reads Mapped Confidently to Exonic Regions | Reads Mapped Confidently to Transcriptome |
| --- | --- | --- | --- | --- | --- | --- |
| Stage2 | 94.19% | 75.11% | 18.73% | 9.80% | 46.58% | 54.97% |

3. Quality metrics of spatial transcriptome data for sample

| sampleID | Number of Cells | Median Genes per Cells | Median UMI Counts per Cells | Total Genes Detected |
| --- | --- | --- | --- | --- |
| Stage2 | 1827 | 2650 | 4253 | 47806 |

Supplementary Table. 3

Summary of read counts, median genes detected, and UMI counts per spot under different resolutions

| SampleID | level | Number of SupSpots | Median UMI Counts per SupSpot | Median Genes per SupSpot | Total Genes Detected |
| --- | --- | --- | --- | --- | --- |
| Stage2 | 18 | 94 | 126128 | 22836 | 48424 |
| Stage2 | 9 | 395 | 29455 | 10968 | 48413 |
| Stage2 | 7 | 673 | 16725 | 7570 | 48332 |
| Stage2 | 6 | 936 | 11849 | 5972 | 48325 |
| Stage2 | 5 | 1394 | 8016 | 4420 | 48326 |
| Stage2 | 4 | 2317 | 4855 | 2959 | 48353 |
| Stage2 | 3 | 4507 | 2469 | 1673 | 48348 |
| Stage2 | 2 | 12181 | 902 | 680 | 48342 |
| Stage2 | 1 | 61714 | 178 | 150 | 48338 |

Supplementary Table. 4

Primer sequences used in this study

| Gene ID | Primer Sequence (5' → 3') | Purpose |
| --- | --- | --- |
| Potri.005G211800/POTOM_021849 | F:ATGGCAGTTGGAAGGATGG;R:TTATATGGTGCCGAGGGTCG | In situ hybridization |
| Potri.002G189300/POTOM_010703 | F:ATGGCAAGCATGACACTTTCA;R:GCTGAAGATTTGAGCACCAGC | In situ hybridization |
| Potri.004G086600/POTOM_017200 | F:ATGGCTTCTTCAATGAGCTTGAAGCTGGCCT;R:TCACTTCACGCTTTTGCAAT | In situ hybridization |
| Potri.002G046400/POTOM_009542 | F:ATGGCTGGAAAAGGAGGAA;R:TCACTCTTTGGTGGTTTTGTTG | In situ hybridization |
| Potri.014G072300/POTOM_047904 | F:ATGGCCTCCATTGATGTTCTTTCAAATCCACT;R:CTAAGTAGTGTTTCTGTGGAAGGCA | In situ hybridization |
| Potri.006G237500/POTOM_024085 | F:ATGATGGCTGTAAGCTCGACG;R:TCACACAAAAGACCAGTTGATGAAC | In situ hybridization |
| Potri.007G068300/POTOM_027161 | F:ATGGCTTCTTTCTTTCATCGTGT;R:ATATGCAAGTGCTGTCTTAACACC | In situ hybridization |
| Potri.006G239700/POTOM_026303 | F:ATGGCAAAAATTGCGTTGG;R:TTAGAAAGGCTCTTCTTCGCTG | In situ hybridization |
| PtoPDF2-pbi121 | F:AGAACACGGGGGACTATGTTCCACGCAAATATGTTTG;R:ACCCCCGGGGATCCTGGCGTTGTCACAAATTACTGC | subcellular localization |
| PtoPDF2-AD | F:GCCATGGAGGCCAGTGAATTCATGTTCCACGCAAATATGTTTG;R:CAGCTCGAGCTCGATGGATCCTCAGGCGTTGTCACAAATTACT | Y1H |
| Pro-PtoMYB38 | F:GAAAAGCTTGAATTCGAGCTCGGCCATCTATCATCAGTCATGTT;R:AGCACATGCCTCGAGGTCGACCGGCTTGATACATTGAAAAAAATT | Y1H |
| Pro-PtoSTK2 | F:GAAAAGCTTGAATTCGAGCTCGCAGGTCTGTGCTCCTTTGT;R:AGCACATGCCTCGAGGTCGACGAGAAGAAGAGAAAACACTGAAACG | Y1H |
| PtoPDF2-SK | F:CGCTCTAGAACTAGTGGATCCATGTTCCACGCAAATATGTTTG;R:GTCGACGGTATCGATAAGCTTTCAGGCGTTGTCACAAATTACT | LUC |
| PtoGAI-SK | F:CGCTCTAGAACTAGTGGATCCATGAAGAGAGATCATCAAGAAACCA;R:GTCGACGGTATCGATAAGCTTTCATTGTTGTGAATCACCAGC | LUC |
| PtoMYB38-LUC | F:GTCGACGGTATCGATAAGCTTGGCCATCTATCATCAGTCATGTT;R:CGCTCTAGAACTAGTGGATCCCGGCTTGATACATTGAAAAAAATT | LUC |
| PtoSTK2-LUC | F:GTCGACGGTATCGATAAGCTTGCAGGTCTGTGCTCCTTTGT：R:CGCTCTAGAACTAGTGGATCCGAGAAGAAGAGAAAACACTGAAACG | LUC |
| PtoMYB38-probe | ACTAAAAAAAGTAAATGTACAAATAAATAG | EMSA |
| PtoSTK2-probe | AAATGTTCTTATAAATGCATGGTTTCGTGC | EMSA |
| PtoPDF2-pSPYNE | F:ggcgcgccactagtggatccATGTTCCACGCAAATATGTTTG;R:gcggtaccctcgaggtcgacGGCGTTGTCACAAATTACTGC | Bifc |
| PtoGAI-pSPYCE | F:ggcgcgccactagtggatccATGAAGAGAGATCATCAAGAAACCA;R:gcggtaccctcgaggtcgacTTGTTGTGAATCACCAGCG | Bifc |
| PtoPDF2-Nluc | F:GAGCTCGGTACCCGGGATCCATGTTCCACGCAAATATGTTTG;R:GCGTACGAGATCTGGTCGACGGCGTTGTCACAAATTACTGC | LCA |
| PtoGAI-Cluc | F:CGGGGCGGTACCCGGGATCCATGAAGAGAGATCATCAAGAAACCA;R:CGAAAGCTCTGCAGGTCGACTCATTGTTGTGAATCACCAGC | LCA |

Note: F, forward primer; R, reverse primer. All primer sequences were designed based on the *P.tomentosa* coding sequences and synthesized by Beijing Ruibo Biotechnology
